# Metadata preservation and stewardship for genomic data is possible, but must happen now

**DOI:** 10.1101/2022.09.12.507034

**Authors:** Eric D. Crandall, Rachel H. Toczydlowski, Libby Liggins, Ann E. Holmes, Maryam Ghoojaei, Michelle R. Gaither, Briana E. Wham, Andrea L. Pritt, Cory Noble, Tanner J. Anderson, Randi L. Barton, Justin T. Berg, Sofia G. Beskid, Alonso Delgado, Emily Farrell, Nan Himmelsbach, Samantha R. Queeno, Thienthanh Trinh, Courtney Weyand, Andrew Bentley, John Deck, Cynthia Riginos, Gideon S. Bradburd, Robert J. Toonen

## Abstract

Genetic diversity within species represents a fundamental yet underappreciated level of biodiversity. Because genetic diversity can indicate species and population resilience to changing climate, its measurement is relevant to many national and global conservation policy targets. Many studies of evolutionary biology, molecular ecology and conservation genetics produce large amounts of genome-scale genetic diversity data for wild populations. While open data policies have ensured an abundance of freely available genomic data stored in the databases of the International Nucleotide Sequence Database Collaboration (INSDC), only about 13% of current accessions have the associated spatial and temporal metadata in INSDC necessary to be reused in monitoring programs, macrogenetic studies, or for acknowledging the sovereignty of nations or Indigenous Peoples. We undertook a “distributed datathon” to quantify the availability of these missing metadata in sources external to the INSDC and to test the hypothesis that these metadata decay with time. We also worked to remediate these missing metadata by extracting them, when present, from associated published papers, online repositories, and/or from direct communication with authors. Starting with 848 programmatically identified candidate datasets (INSDC BioProjects), we manually determined that 561 contained samples from wild populations. We successfully restored spatiotemporal metadata (locality name and/or geospatial coordinates and collection year) for 78% of these 561 datasets (N = 440 BioProjects comprising 45,105 individuals or BioSamples from 762 species in 17 phyla). We also quantified the availability of 33 additional categories of metadata in sources external to the INSDC. Information about associated publications and the type of habitat from which the samples were taken was the most easily found; information about sampling permits was the most challenging to locate. Looking at papers and online repositories was much more fruitful than contacting authors, who only replied to our email requests 45% of the time. Overall, 23% of our email queries to authors discovered useful metadata. Importantly, we found that the probability of retrieving spatiotemporal metadata declines significantly with the age of the dataset, with a 13.5% yearly decrease for metadata located in published papers or online repositories and up to a 22% yearly decrease for metadata that were only available from authors. This observable metadata decay, mirrored in studies of other types of biological data, should motivate swift updates to data sharing policies and researcher practices to ensure that the valuable context provided by metadata is not lost forever.

## Introduction

Genetic diversity is the foundational layer of biodiversity. Just as ecosystem health and resilience depends on the diversity of its component species, the health and resilience of each species depends on its genomic diversity (Clark, 2010; Reusch, Ehlers, Hämmerli, & Worm, 2005). Without genetic diversity in the form of standing allelic variation, populations and species cannot adapt to a rapidly changing climate and other anthropogenically-induced stresses (Blanchet et al., 2020; Raffard, Santoul, Cucherousset, & Blanchet, 2019). Local or global extinctions of species in turn threaten the ecosystems upon which the quality of human lives depend (Brauman et al., 2020; Des Roches, Pendleton, Shapiro, & Palkovacs, 2021). Concerningly, genetic diversity, like all levels of biodiversity, is declining rapidly during the Anthropocene across the tree of life (Exposito-Alonso et al., 2022; Leigh, Hendry, Vázquez-Domínguez, & Friesen, 2019; Miraldo et al., 2016; Pinsky & Palumbi, 2014).

Recognizing the vital importance of biodiversity to human well-being and the future of our planet, several international agreements strongly encourage the monitoring and conservation of genetic diversity in both wild and domesticated species. Foremost among these are the United Nations Sustainable Development Goal 2.5 and the international Convention on Biological Diversity (CBD) treaty, which explicitly acknowledge the importance of monitoring and conserving any component of biological diversity (including genetic diversity) that may have “actual or potential use or value for humanity.” Moreover, the CBD’s article 15 and attendant Nagoya Protocol codify procedures to ensure the sharing of benefits arising from genetic resources (such as digital sequence information; DSI) discovered or accessed within a nation’s sovereign borders. The subsequent Strategic Plan for Biodiversity 2011-2020 laid out the 20 Aichi Biodiversity Targets, including target 13, which aims to maintain the “genetic diversity of cultivated plants and farmed and domesticated animals and of wild relatives, including other socio-economically as well as culturally valuable species.” Now, even as we are facing shortfalls on all 20 of the Aichi Biodiversity Targets (CBD, 2020; Hoban et al., 2021), there have been calls to broaden genetic diversity targets to include *all* extant species in the New Post-2020 Global Biodiversity Framework, in what has been described as a “moonshot for biology” (Hoban et al., 2020; Laikre et al., 2020; Lewin et al., 2018).

Over the last decade, advances in DNA sequencing technology have enabled the generation of genome-scale datasets of ever larger numbers of individuals, drawn from a growing variety of species (Allendorf, 2017; Hendricks et al., 2018). Researchers are now able to genotype thousands of genomic loci or sequence whole genomes from non-model species for which they have no prior genetic resources (Lou, Jacobs, Wilder, & Therkildsen, 2021; Willette et al., 2014). The shift from genetic to genomic-scale datasets is catalyzing novel conservation insights including: the detection of inbreeding depression (e.g. Kardos, Taylor, Ellegren, Luikart, & Allendorf, 2016), the discovery of subtle, previously undetectable population structure (e.g. Cheng, Gold, Rodriguez, & Barber, 2021; Gaither et al., 2018), reconstruction of demographic histories (Prada et al., 2016), the precise identification of distant pedigree relationships (e.g. Baetscher et al., 2019), uncovering cryptic species (e.g. Quattrini et al., 2019), clues about the genomic basis of local adaptation (e.g. Wilder, Palumbi, Conover, & Therkildsen, 2020) and important traits such as nutritional components (e.g. Kumar et al., 2021). Accordingly, the DSI derived in these studies is highly valued as a resource equivalent to biobanks, providing essential information for conservation (Hoban et al., 2022) as well as ensuring future food security (Castañeda-Álvarez et al., 2016; Halewood et al., 2018).

Genomic datasets record the genetic diversity of a species at a particular time and location, providing a benchmark for how populations are responding to human drivers of changing environmental conditions, cultivation, and land and sea use, as well as measuring indicators of progress toward conservation targets and goals (Hoban et al., 2022, 2020) and the genetic resources available for future cultivation or domestication (Halewood et al., 2018). However, genomic datasets can only be useful for monitoring global genetic biodiversity and the sustainable human use of genetic diversity (including benefit-sharing, Cowell et al., 2022) when archived publicly with accompanying metadata about the spatiotemporal, environmental and methodological context of the sequenced sample (Riginos et al., 2020; Scholz et al., 2022; Schriml et al., 2020).

The genetics community has long championed open data publication with the foundational databases of the International Nucleotide Sequence Database Collaboration (INSDC; Cochrane, Karsch-Mizrachi, Takagi, & INSDC, 2016) forming in the early 1980’s. In 2009, the INSDC launched the Sequence Read Archive as a repository dedicated to second-generation sequence data. It has since grown exponentially to include over 600 terabytes of freely-available DNA sequence data from over 16,700 wild and domesticated eukaryotic species as of 2021 (Toczydlowski et al., 2021). Around the same time, the MIxS metadata standards (Field et al., 2008; Yilmaz et al., 2011) were defined to inform the minimum information about *what* (detailed taxonomy), *where* (GPS coordinates and habitat), *when* (collection date), *how* (sampling and sequencing protocols) and *by whom* a genetic sample was collected. Enabled by the INSDC infrastructure and encouraged by the Joint Data Archiving Policy (JDAP; http://datadryad.org/pages/jdap) implemented by top journals in 2011, the proportion of papers providing open access to their genetic data increased dramatically (Pope, Liggins, Keyse, Carvalho, & Riginos, 2015). However, the inclusion of accompanying metadata crucial for the reuse of these data for genetic diversity monitoring, macrogenetic studies, or identifying their provenance within national boundaries or the lands and waters of Indigenous Peoples, has lagged behind (Pope et al., 2015; Toczydlowski et al., 2021). As of 2021, out of over 300,000 SRA BioSamples that are potentially relevant to global genetic biodiversity, only ∼13% had metadata indicating both the time and precise location from which they were sampled (Toczydlowski et al., 2021).

In a timely and welcome update to their policy, INSDC now intends to extend their minimum metadata requirements to include collection date and country of origin (https://www.insdc.org/spatio-temporal-annotation-policy-18-11-2021). Although ‘country’ is legislatively aligned with the Nagoya Protocol, it is not spatially aligned with the lands and waters of Indigenous Peoples (e.g. https://native-land.ca/) and does not provide adequate spatial resolution for conservation monitoring. Moreover, this policy and infrastructure change will take time to implement (anticipated to be end of 2022), meaning that much of the genomic data collated over the last ∼12 years for past and present populations, of immeasurable value to understanding and monitoring the biodiversity crisis, are not Findable, Accessible, Interoperable or Reusable (FAIR; Wilkinson et al., 2016). This absence of appropriate spatio-temporal metadata represents the effective loss of tens to hundreds of millions of dollars of research effort for most future purposes (Schriml et al., 2020; Toczydlowski et al., 2021), rendering associated genetic data invisible to government ministries and non-governmental organizations tasked with protecting the world’s natural environment (Laikre, 2010; Laikre et al., 2020). Moreover, without spatiotemporal provenance of genomic data enabling connection to the lands and waters of Indigenous Peoples, these peoples will potentially lose out on benefits (e.g. capacity development, food security, biomedical advances) arising from genomic information originating within their territories (Liggins, Hudson, & Anderson, 2021; Marden et al., 2021; McCartney et al., 2022; Scholz et al., 2022). There is urgency in addressing this metadata gap: previous studies of morphological (Vines et al., 2014) and genetic (Pope et al., 2015) data suggest that the probability of existing metadata ever being linked to the genomic data significantly decreases over time.

In the summer of 2020, we convened a distributed remote datathon to (1) assess the availability of metadata outside of the INSDC, (2) recover and curate metadata missing in INSDC from external sources (i.e. published research papers, other online repositories, or the authors themselves), and (3) extend our initial report on the metadata gap (Toczydlowski et al., 2021) to investigate how the recoverability of these metadata is affected by dataset age and to document shortfalls and costs of our remedial efforts. In our datathon, 14 graduate students from the USA and 12 professional academics (authors on this paper) across 3 countries worked together via Zoom, Slack and Google Sheets as “metadata curators” to establish and execute curation protocols and infill missing metadata. Collectively, we searched for metadata external to the INSDC (e.g. associated scientific publications, Dryad, museum collections) for 848 genomic datasets (INSDC BioProjects) representing 94,416 individual samples (BioSamples). The BioSamples and associated genetic sequence data in these projects were selected as they were missing at least latitude and longitude metadata in the INSDC. Our findings underscore the importance of appropriate and immediate metadata archival going forward. We provide guidance based on our collective experience gained over the datathon on practices to retain crucial metadata.

## Methods

### Identifying BioProjects for metadata curation

Our datathon followed a workflow illustrated in Figure 1. We first searched the INSDC to identify datasets that were potentially relevant to monitoring genetic diversity of wild populations (including wild relatives of domesticated species) but were lacking critical metadata about the latitude and longitude of the sampling location (as described in Toczydlowski et al., 2021). On November 7, 2019, we searched the INSDC BioProject database using the *rentrez* R package (Winter, 2017) and custom R scripts to locate datasets that contained genomic-level DNA sequence data from eukaryotic species, excluding human and metagenomic datasets. We then downloaded all metadata associated with these BioProjects from the INSDC PubMed, Taxonomy, BioSample and SRA databases. We further filtered the BioProjects to remove BioSamples (sequenced individuals) from species whose population dynamics and evolution are largely governed by humans: pathogens and their vectors, model organisms, and domesticated species. We used custom lists for each category of non-wild organisms (Supplementary Materials S1). These filters helped to target our efforts at recovering metadata for the datasets most likely to be of relevance to genetic diversity monitoring of wild populations. Additional details about the search and filtering steps can be found in Supplemental Materials of Toczydlowski et al. 2021. These filtering steps yielded 848 BioProjects that entered our datathon curation pipeline. These BioProjects contained at least 5 individuals that lacked geospatial coordinates in the INSDC and were potentially sampled from wild populations.

**Figure 1.**
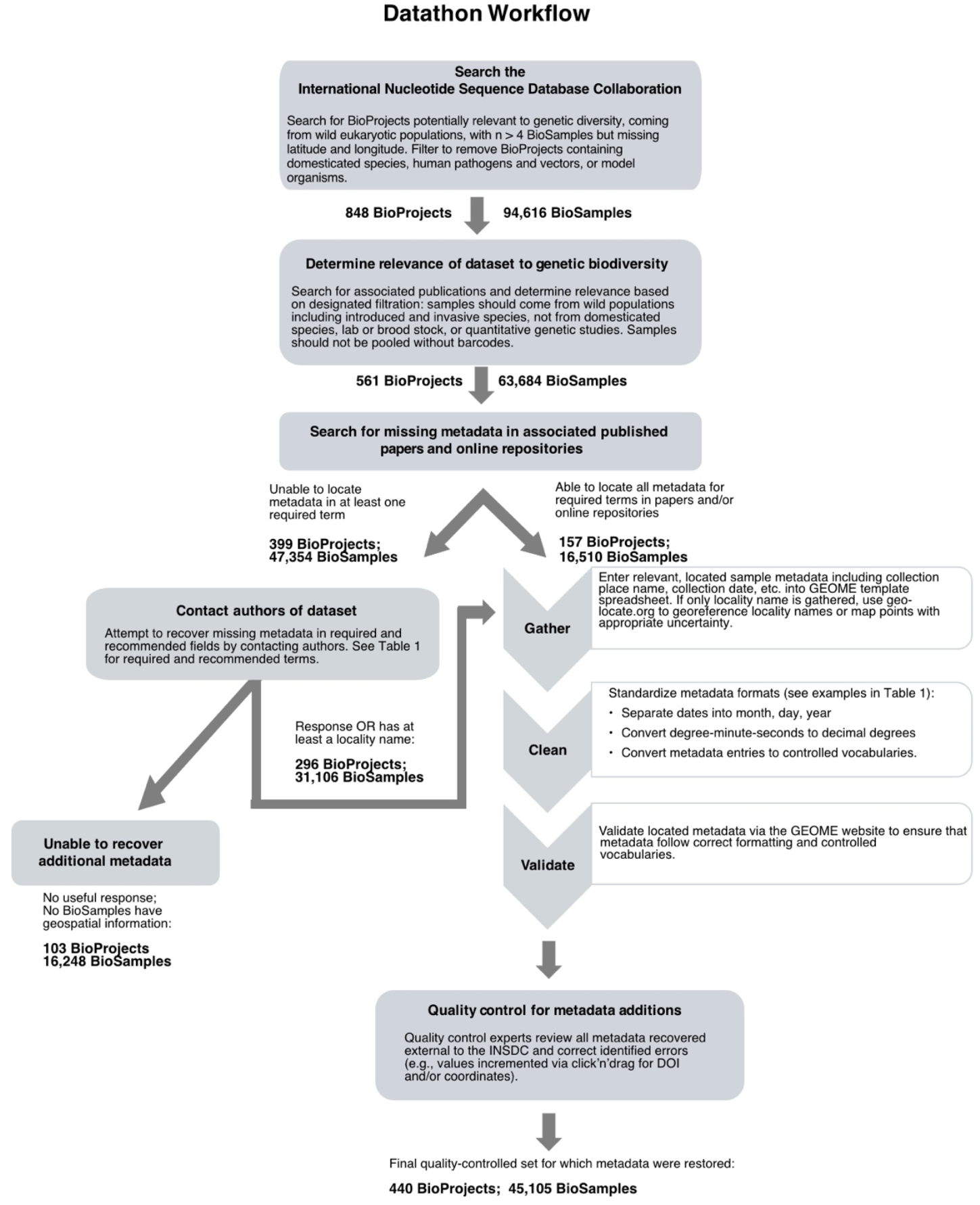
Datathon workflow. The number of BioProjects and BioSamples remaining after each step are given below the step.

We built a blank template to enter metadata into that we located external to the INSDC using the Genomic Observatories Metadatabase (GEOME; Deck et al., 2017; Riginos et al., 2020) This template included terms that described collection date and location, habitat etc. Table 1 gives definitions of 16 required and recommended metadata terms, while Supplemental Materials S2 provides an example template with all 36 metadata terms and their definitions and controlled vocabularies. We then programmatically filled this template for each of the 848 BioProjects in our pipeline with the metadata already present in the INSDC using custom R scripts.

**Table 1:**
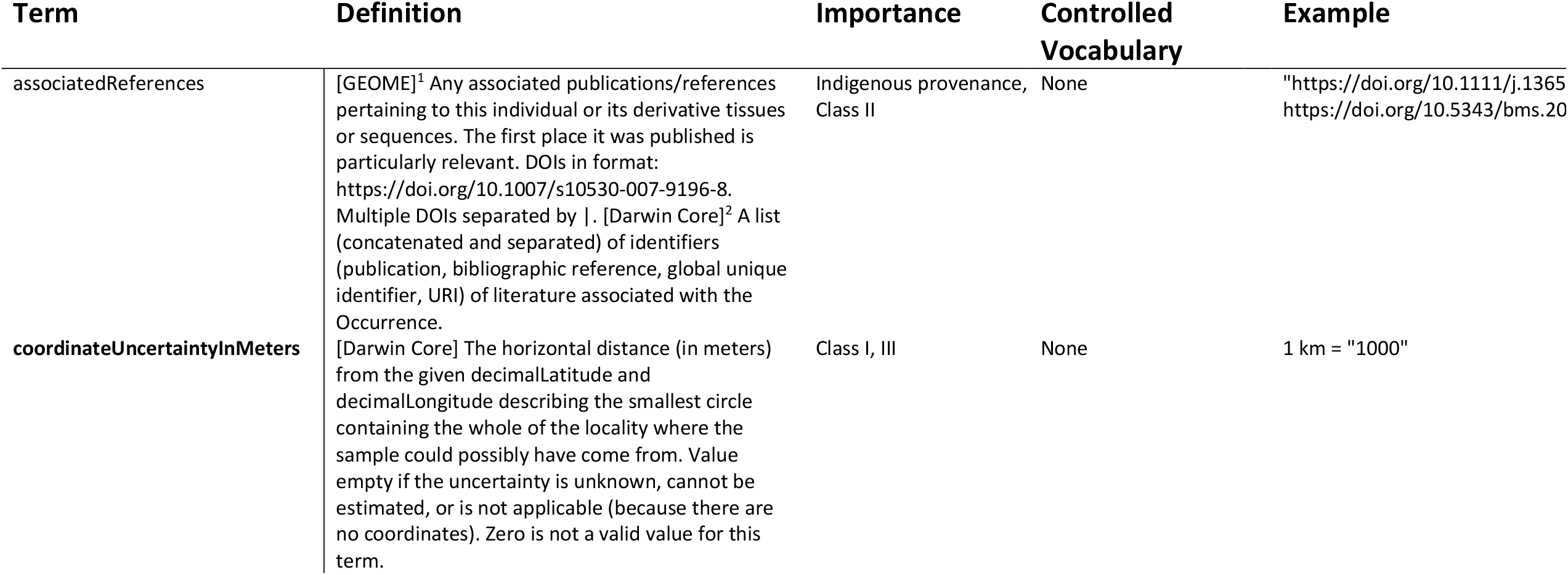
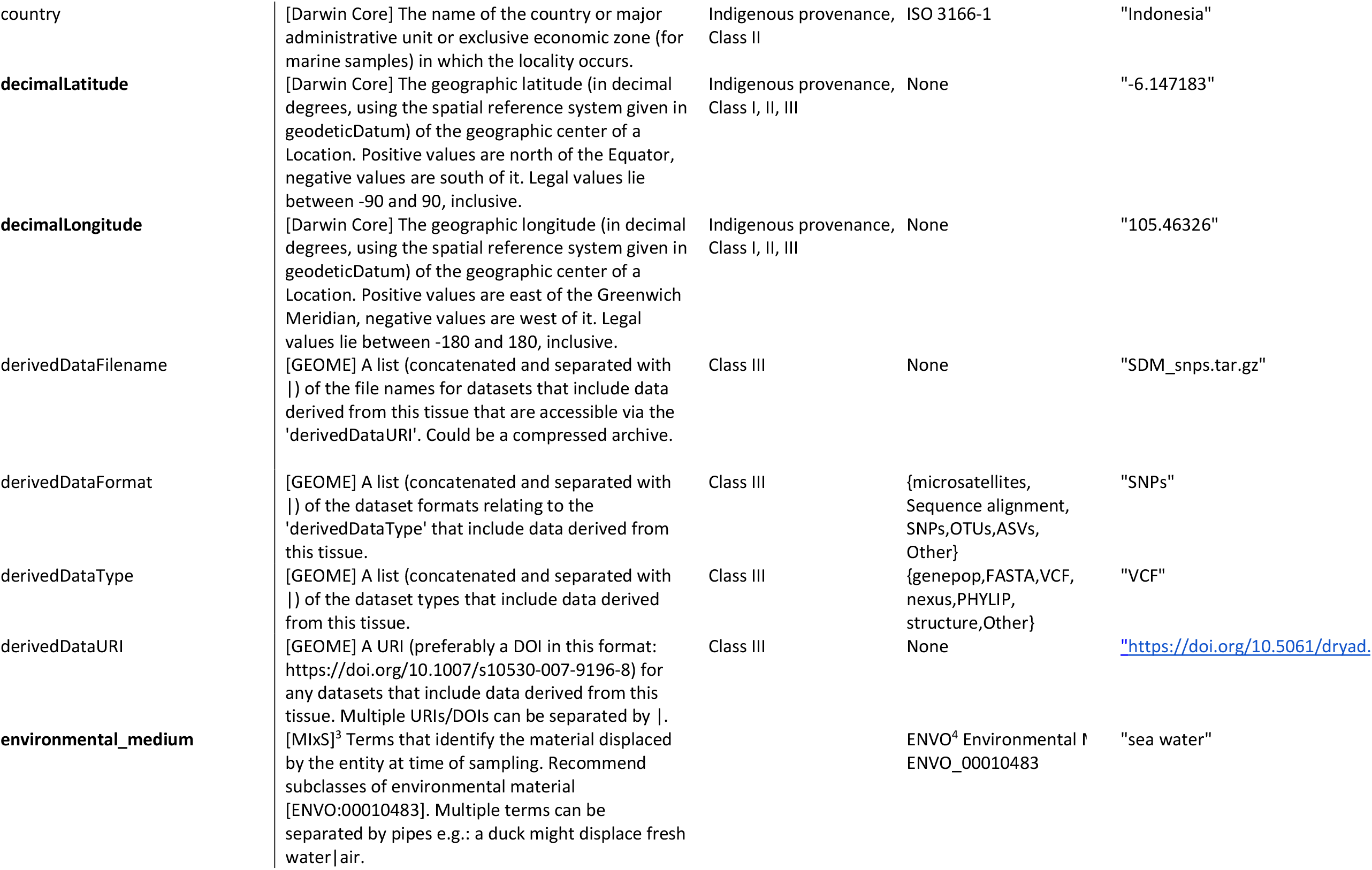
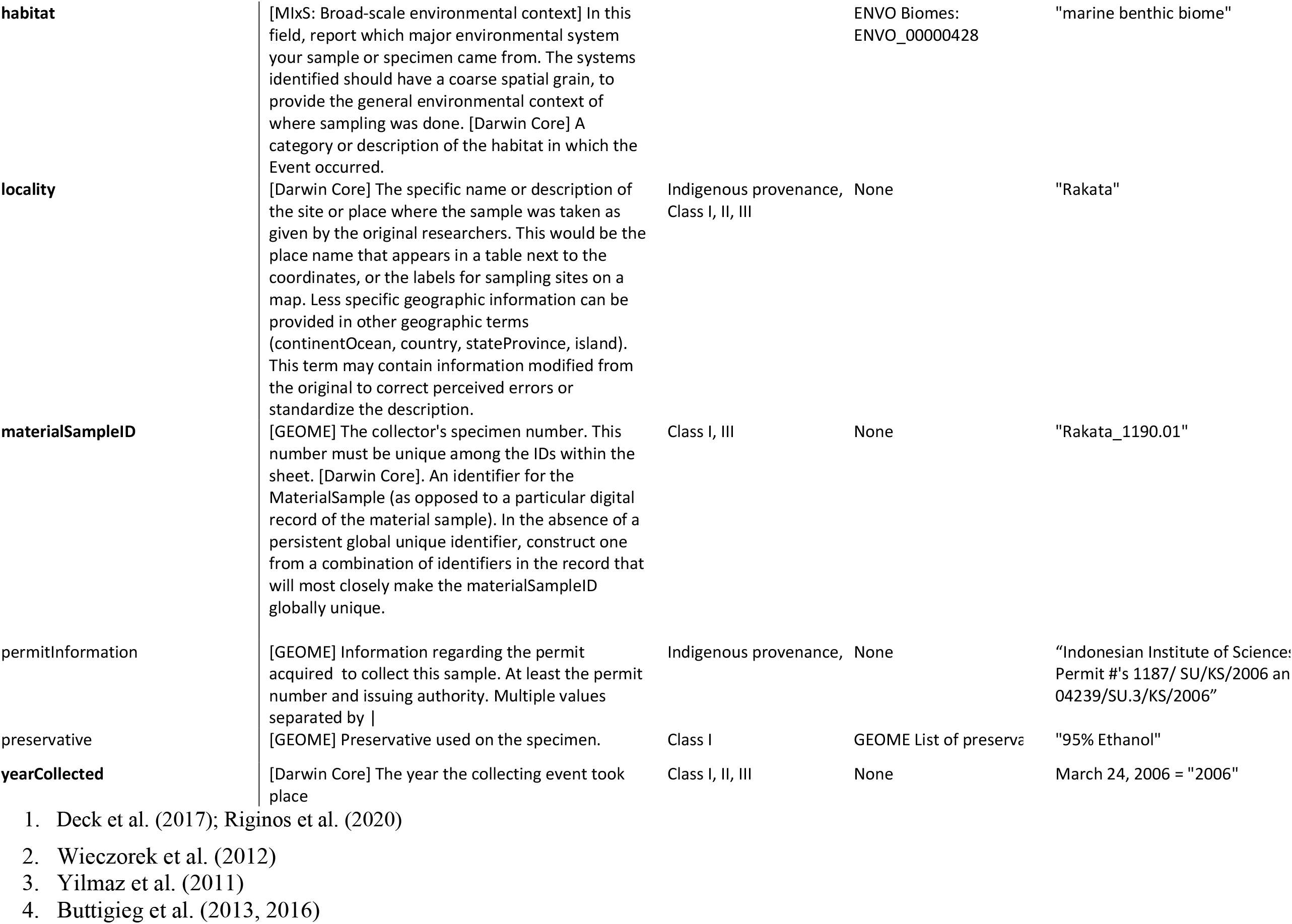
Alphabetized list of required (in bold) and recommended metadata terms for individual organisms and/or derived tissues or DNA sequences included in the datathon. Square brackets in the definition column denote the metadata standard from which the definition comes. Terms with multiple definitions are in order of decreasing specificity. The importance column indicates which terms support the identification of Indigenous provenance and can therefore inform Access and Benefit-Sharing (ABS), and those that can support sample or Digital Sequence Information (DSI) re-use in conservation, according to the study approach definitions of Leigh et al. (2021). Class I studies generate new sequence data, requiring precise information regarding the spatiotemporal context of the collected sample, a unique materialSampleID, as well as the preservative the tissue is held in; Class II studies compile genetic diversity values from published studies, generally requiring less precise spatiotemporal information, but this needs to be associated with a publication (associated References); Class III studies re-analyse digital sequence information, or derived genetic data, requiring precise spatiotemporal information, and a unique materialSampleID. Depending on the objective of re-use, habitat and environmental_medium may also be important for sample/DSI re-use in conservation. Controlled vocabularies refer to standardized lists of acceptable entries, often defined by a standards organization.

### Recovering and curating metadata external to the INSDC

Metadata curators were each randomly assigned a set of BioProjects. Curators followed a standard protocol (Figure 1; Supplemental Materials S3) to locate and enter associated metadata for samples in each BioProject that were missing in the INSDC but reported in external sources (e.g. associated published scientific papers). Briefly, we first searched for associated publications by googling the BioProject PRJ accession number and/or key words associated with the BioProject (e.g. author, affiliation, funding information, species, and locality names). While the INSDC includes terms to capture associated publications, these were only populated for 7% of the 848 BioProjects that we examined. We then skimmed each paper to determine if the BioProject contained sequence data from wild individuals. BioProjects comprising over 50% BioSamples from non-wild individuals (brood stock, laboratory stock, domesticated species, and some seed banks) were marked as “not relevant” and removed from further curation. Non-wild BioSamples that were a minority in their BioProject were designated as such under the *establishmentMeans* metadata term, and further metadata recovery was not prioritized for these samples. BioProjects that pooled their genetic samples without barcodes (precluding estimates of genetic diversity at the level of individual) were also marked as “not relevant” and removed from the curation pipeline. Once relevance to our efforts was determined, curators looked in figures, tables, and supplemental information of associated publications and/or linked online repositories (e.g. Data Dryad, Github, Zenodo, and databases of biocollections such as museums and herbaria) for sample-level metadata from 36 metadata terms (most defined as Darwin Core terms; Wieczorek et al., 2012). When necessary, curators converted metadata to standard formats (e.g. degree minutes seconds data were converted to decimal degrees), and then added metadata into the pre-generated GEOME template for that BioProject. After performing quality control, these metadata could then be easily uploaded to GEOME and potentially cross-walked into the appropriate INSDC databases.

After adding all metadata that could be gleaned from the associated paper(s) into the GEOME templates, curators made a structured comment on a master spreadsheet (Supplemental Materials S4) indicating whether metadata for each of the required and recommended terms were absent for all BioSamples (“none”), present for less than 50% of BioSamples (“some”), present for greater than 50% of BioSamples (“most”), or iv) present for all BioSamples (“all”). If the paper was missing information from one of six or seven “required” terms (georeference-able *locality* OR [*decimalLatitude* AND *decimalLongitude*], *coordinateUncertaintyInMeters, georeferenceProtocol, habitat, environmentalMedium, yearCollected*), the curator flagged the BioProject to initiate author contact. An additional nine metadata terms were considered “recommended”: missing metadata in these fields alone did not trigger an author contact but curators and authors were asked to populate these fields as completely as possible. These recommended terms included *country, establishmentMeans, permitInformation, associated References, preservative*, and 4 termss that tracked genetic data derived from the raw reads such as SNP genotypes or sequence alignments. Progress and notes at each curation step were tracked as meta-metadata on the master spreadsheet.

After a quality-control step to ensure that author names and email addresses found in papers were input correctly, corresponding authors of the paper were contacted by email (see Supplemental Material S5 for email text) using the Yet Another Mail Merge add-on for Google Sheets (yamm.com). Using YAMM allowed us to track email receipts and responses. If an email was undeliverable, we used our best efforts to locate an alternate email. We were able to successfully deliver email queries for all 351 of 492 relevant BioProjects that met the criteria for author contact. About two weeks after sending the initial email, curators sent reminder emails to unresponsive authors at least once and at most twice. This process emulated the efforts of a reasonably persistent researcher to obtain metadata important to their research.

Once curators had gathered and entered all available metadata into the GEOME template for a BioProject (from online sources external to the INSDC and/or directly from authors), curators validated the metadata at the GEOME website (geome-db.org). This programmatic GEOME validation ensured that metadata in each field were correctly formatted and within controlled vocabularies, e.g. for *country* and *habitat* (see lists tab of Supplemental Materials S3 for all controlled vocabularies). Following validation, MRG, BEW, AP and EDC performed a final quality-control check (QC; steps described in Supplemental Material S1). Filled and QCed GEOME templates for each BioProject will be uploaded to the GEOME database.

### Investigating metadata decay

We investigated the effect of BioProject age on the probability that we were able to recover metadata information for 11 metadata categories. Previous investigations of metadata have indicated rapid decay when data are not publicly archived (Roche et al., 2014; Vines et al., 2014, 2013). We used Bayesian logistic regression to fit four distinct models to investigate the relationships between BioProject age (number of days between publication in the INSDC and November 7, 2019) and: A) the probability that metadata could be retrieved from INSDC, associated published papers, and/or repositories, B) the probability that we received an author response for the 351 BioProjects that triggered an author contact via email, C) the probability that authors provided any metadata, given that they responded and D) the probability that authors provided metadata for a majority of samples, given that they responded.

Information about the collection date and location of a sample are the most critical pieces of metadata required to identify the Indigenous provenance of genomic data and make genomic sequence data repurposable for monitoring and fundamental data synthesis research, so we focused our investigations on these two categories; we refer to the aggregate as spatiotemporal metadata. We defined a BioProject as having spatiotemporal data if collection dates, and latitudes and longitudes and/or locality were present for at least 50% of the BioSamples that it contained. In model C, we counted a gain in collection year, or place name, or latitude/longitude for any number of BioSamples as recovery of metadata. In model D, we only counted increases in metadata where BioProjects had incomplete spatiotemporal metadata for > 50% of its BioSamples and then had spatiotemporal metadata present for > 50% of BioSamples after contacting authors. That is, model C assessed the probability of recovering any metadata external to the INSDC, and model D assessed the probability of recovering metadata for the majority of samples. In a supplemental analysis, we investigated how metadata within individual spatiotemporal terms and other important metadata terms (i.e. required and recommended terms, Table 1) decayed, including: *decimalLongitude and decimalLatitude, collectionYear, country, locality, habitat, environmentalMedium, materialSampleID, permitInformation, associatedReferences* (publication DOI), *preservative*, and derived genetic data terms (Supplemental Materials S6).

We conducted all statistical analyses at the level of BioProject (as opposed to BioSamples or genomic sequences), because presence/absence of metadata for BioSamples within a given BioProject was highly correlated (Toczydlowski et al., 2021). Curators also tracked metadata recovery efforts at the level of BioProject for convenience, and we contacted authors about entire BioProjects rather than individual BioSamples. In each set of models, we removed BioProjects that already had complete metadata in the category of interest, and therefore could not gain any more.

We analyzed the effect of BioProject age on our response variables (model A: successful metadata retrieval; model B: author response; model C: author provided any metadata conditional on a response to our query; model D: author provided metadata for the majority of samples conditional on a response to our query) using generalized linear models. In each analysis, we modeled our response variable as a Bernoulli-distributed variable with a probability of success that was a linear function of our predictor variable: BioProject age. In each analysis, the parameters of our model were a global mean probability of success and an effect size of BioProject age on probability of success for that response variable. Analyses used the canonical inverse-logit inverse link function. In mathematical notation, our model was:

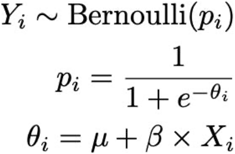

Where *Y*_*i*_ is the *i*th outcome (response variable), *p*_*i*_ is the probability of successfully observing that outcome, μ is the global mean probability of success, and β is the effect of BioProject age on the transformed probability of success for that outcome (θ_i_). We had no strong prior beliefs about the effect of BioProject age on success in each of the four analyses we ran; to reflect these beliefs, the priors we placed on our parameters were: β ∼ N(0,10); μ ∼ N(0,10). All statistical analyses were performed using Rstan version 2.21.2 [50] running 4 independent chains for 2,000 iterations, thinning to sample only every 4th iteration to reduce autocorrelation, and discarding the first 1,000 iterations as burn-in. To assess the significance of the effect of BioProject age on success of each outcome, we determined whether the 95% equal-tailed credible interval of the marginal distribution on β contained 0; if it did, the effect of BioProject age was deemed not significant.

## Results

We identified 848 INSDC BioProjects, representing 94,416 BioSamples from individual eukaryotic organisms that lacked geospatial coordinates and had at least five putatively wild individuals as determined by our filters. Curators were able to locate associated published scientific papers for 741 of these 848 BioProjects. Reading these papers revealed 561 BioProjects with a majority of relevant, truly wild individuals, comprising 63,684 individuals from 873 species. After scouring associated published papers for metadata and contacting authors, a total of 440 BioProjects with 45,105 BioSamples from 762 species in 17 eukaryotic phyla (Figure 2A) had geospatial data (either coordinates or a locality name) and were passed through quality control for eventual upload to GEOME. BioSamples that passed through the datathon came from all continents and all major oceans (Figure 2B-D).

**Figure 2.**
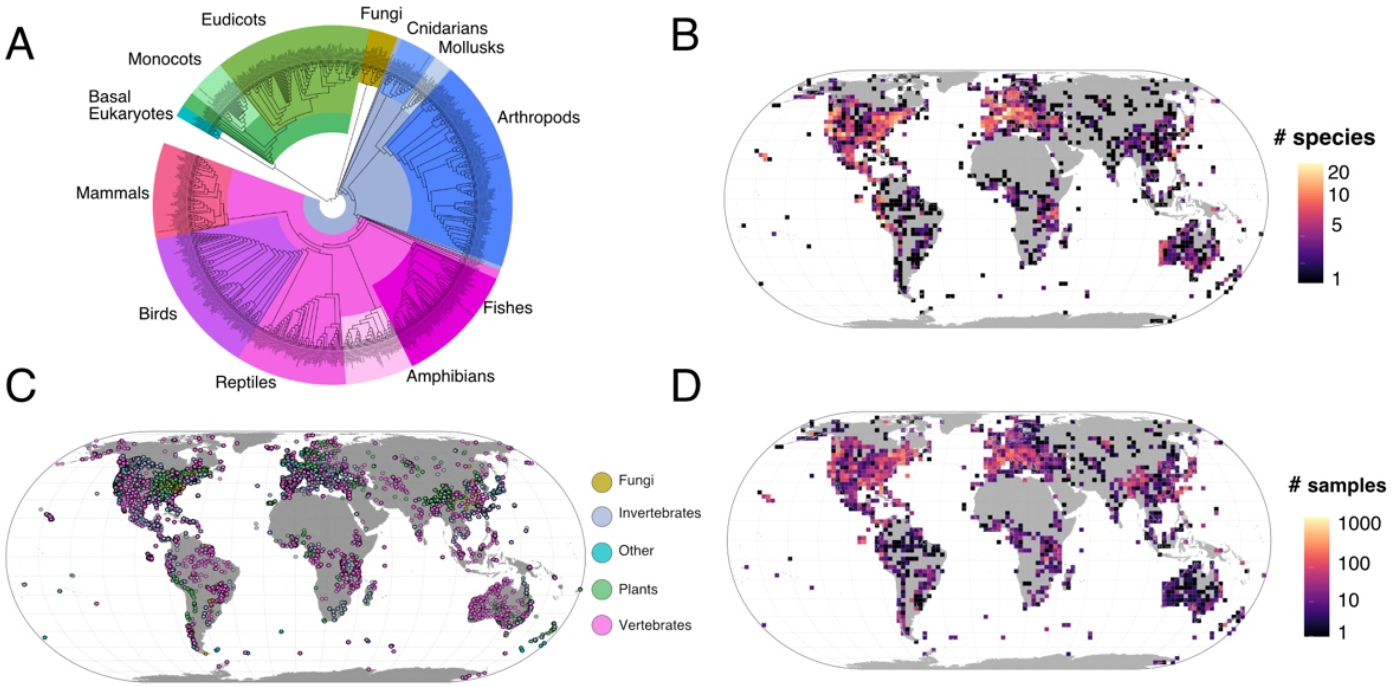
The taxonomic and geographic scope of the datathon. A) A cladogram of 719 of the 762 species from BioProjects that passed through the final quality control step. This is a subtree of the Open Tree of Life (Hinchcliff et al. 2015) generated with the rotl package for R (Michonneau et al. 2016) and visualized with iTOL software (itol.embl.de; Letunic and Bork 2021). B) Heatmap of species for BioSamples for which spatial coordinates were recovered by the datathon. C) Map showing the geographic distribution of broad taxonomic categories of these BioSamples. D) Heatmap of these BioSamples.

For the subset of BioProjects that we focused on (those that were missing latitude and longitude), datathon curators were able to recover metadata for a majority of BioSamples in a BioProject as follows (depicted in Figure 3). For geospatial coordinates, nearly 60% could be found in an associated publication or online repository. While nearly 30% of these BioProjects did already contain information about collection year in the INSDC, curators were only able to recover an additional 21% from papers or online repositories. Datathon curators recovered metadata regarding habitat, environmental medium (the media displaced by the sampled organism) and publication DOI for over 80% of BioProjects from published papers and their supplemental information. Additional large gains in BioProjects were made from online sources external to the INSDC for locality (48.8%), and country name (39.8%). Notably, permit information was the least available of any of the metadata categories that we explored. There is no permit metadata term in the INSDC and curators found permit information in papers for only 21% of BioProjects.

**Figure 3.**
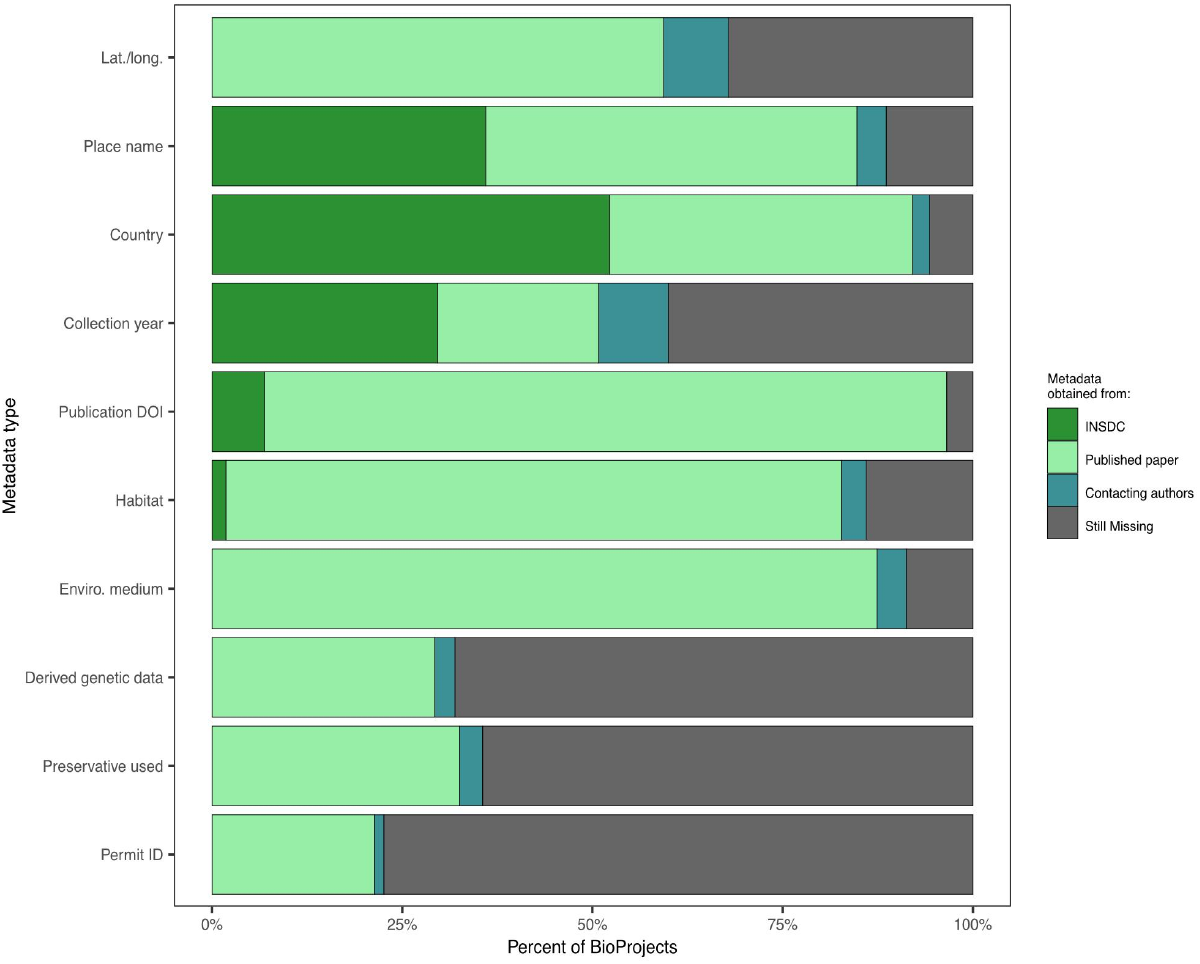
Stacked bars showing the percent of BioProjects for which metadata were found from each of three sources, across 10 priority metadata categories.

Contacting authors yielded comparatively less metadata than our search of papers and supplemental information, although it should be noted that this step was secondary to looking in papers and online. Out of 351 author contact attempts, we received 158 responses (45% response rate). Of the 158 responses, 80 (51%) provided at least some missing metadata, yielding an overall “useful author response rate” of 23%. Through contacting authors, we were able to recover collection year metadata for an additional 9% of BioProjects, and geospatial coordinates for an additional 8.5% percent of BioProjects. Gains in other metadata categories were all less than 5%, with permit information showing only a 1.2% increase from authors.

The age (time since deposition into the INSDC) of the BioProject had a strong effect on whether metadata could be recovered. After searching for metadata within the INSDC and within published papers, we found that spatiotemporal metadata (defined as year AND geospatial coordinates OR locality), had a mean odds ratio of 0.865 (95% highest posterior density credible interval [HPD CI]: 0.775 - 0.964; Figure 4A). This indicates that for every year after a BioProject is published to the SRA, there is about a 13.5% decrease (HPD CI: 3.6% - 22.5%) in the probability that its metadata can be found in the SRA, in papers or elsewhere online. On the other hand, there was a strong positive effect of BioProject age on whether an attempt to contact the authors was answered, with a 25.5% increase in the probability of a reply of any kind for every year after SRA publication (mean odds ratio of 1.255; 95% HPD CI: 1.120 - 1.412; Figure 4B). In other words, we were more likely to get an email response for older datasets. However, given a response, the probability that authors furnished any amount of metadata for year OR coordinates OR locality decreased with BioProject age by 21% per year (odds ratio 0.810; 95% HPD CI: 0.680 - 0.949; Figure 4C). Similarly, the probability that the authors provided metadata for year AND coordinates OR locality for a majority of BioSamples decreased by 22% per year (odds ratio: 0.819; 95% HPD CI: 0.671 - 0.994; Figure 4D).

**Figure 4.**
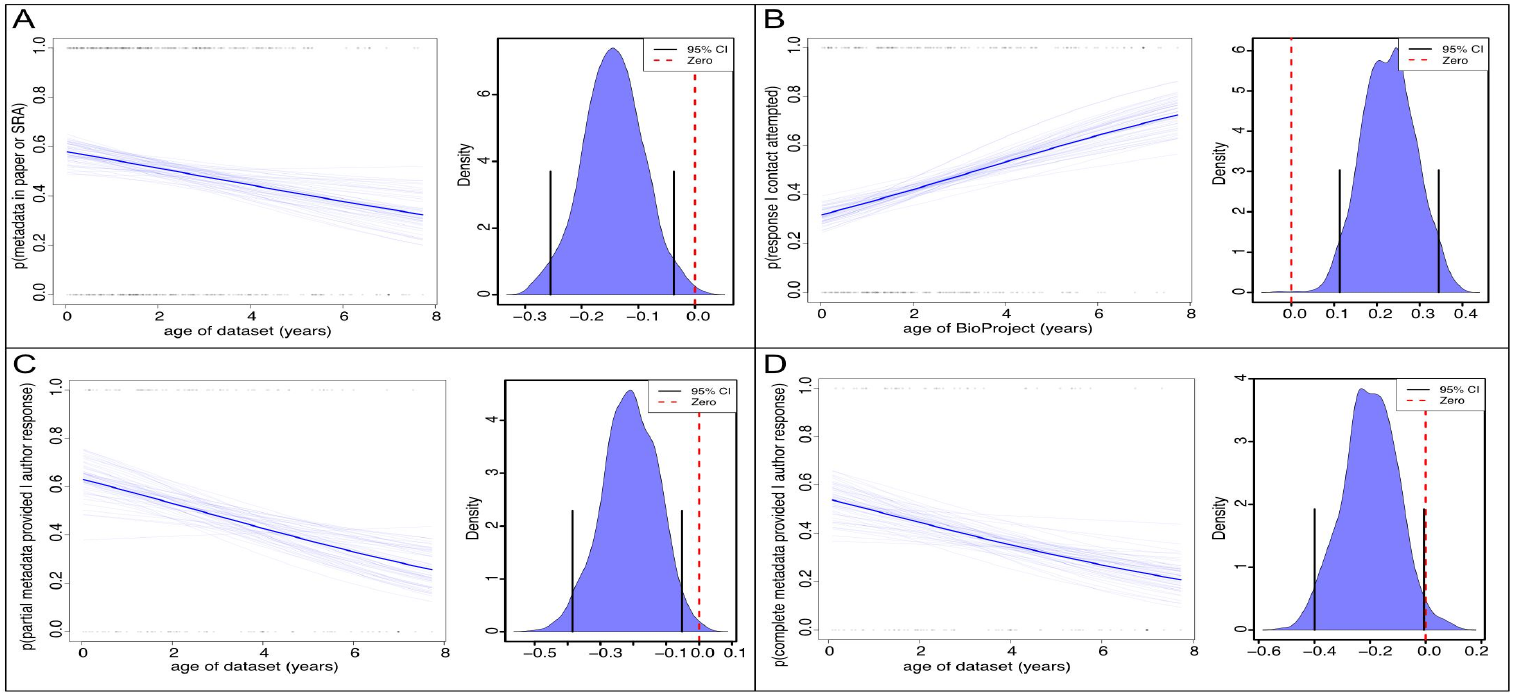
The effect of BioProject age on the probability of recovering spatiotemporal metadata. Density plots depict posterior distribution for log(Odds Ratio), with black lines showing 95% highest posterior density (HPD). (A) Probability that metadata were found in the INSDC or in the INSDC or associated papers and repositories. (B) Probability of receiving a reply from BioProject authors to our contact email. (C) Probability of receiving any amount of additional metadata for year OR coordinates OR locality. (D) Probability of receiving metadata for year AND coordinates OR locality for a majority of BioSamples. 95% HPD intervals exclude 0 in all panels.

Figures for Bayesian logistic regressions of BioProject age on other metadata categories can be found in Supplemental Materials S6 (figures) and S7 (tables of β values). In accordance with the results for spatiotemporal metadata, supplementary analyses indicated that metadata for collection year (posterior mean slope = -0.133, 95% Credible Interval: -0.233 - -0.034; Table S1, Figure S6-11A) and preservative used (posterior mean slope = -0.111, 95% HPD: -0.218 - - 0.009; Figure S6-9A) were significantly less likely to be recovered from INSDC, publications, and/or online repositories with increasing age of a BioProject. Furthermore, and as with spatiotemporal metadata, the probability that responding authors provided additional metadata for georeferences (decimalLatitude and decimalLongitude; posterior mean slope = -0.151, 95% Credible Interval: -0.386 – -0.05; Figure S6-2C), collection year (posterior mean slope = -0.174, 95% Credible Interval: -0.363 – 0.000; Figure S6-11C), and preservative used (posterior mean slope = -0.438, 95% Credible Interval: -0.873 – -0.081; Figure S6-9C) was significantly greater for younger BioProjects. The provisioning of permit information followed this same trend (although marginally insignificant, posterior mean slope = -0.555, 95% Credible Interval: -1.31 - 0.003; Figure S6-5C), suggesting these metadata are relatively available within the personal data management system of authors.

Concerningly and counter to our result for spatiotemporal metadata, supplementary analyses indicated that metadata for habitat (Table S1, posterior mean slope = 0.141, 95% Credible Interval: 0.006 - 0.285; Figure S6-6C) and environmental medium (posterior mean slope = 0.176, 95% Credible Interval: 0.016 - 0.355; Figure S6-5C) were less frequently recovered from INSDC, publications and/or repositories for younger BioProjects. The reasons for these results are unclear, but may indicate a decline in author attention and appreciation of organismal natural history. Retrieval of these metadata through author contact had no relationship with BioProject age.

## Discussion

With this distributed datathon, we have demonstrated that crucial metadata can be restored for many genomic investigations of wild organisms. However, our analyses show that metadata are more difficult to recover the longer we wait, and many are locked in non-standard formats. Because the great majority of publicly available genomic datasets lack important metadata (Toczydlowski et al., 2021), they are not findable, accessible, interoperable nor reusable (FAIR; Wilkinson et al., 2016). Only genomic data that are FAIR will allow systematic monitoring of the fundamental layer of biodiversity (Hoban et al., 2021), and enable assertions regarding provenance for informing CBD Nagoya Protocol obligations. Our results illustrate that: (1) metadata availability is dependent on type: location, publication and habitat metadata are much more available or inferable than metadata about permits and preservatives; (2) with considerable time and paid effort, it is possible to recover some of these important metadata from the non-standardized and non-machine-readable formats in which they are currently being stored; and (3) while metadata archival practices may be improving incrementally, genomic metadata are subject to the same decay processes demonstrated for other types of scientific data (Pope et al., 2015; Vines et al., 2014).

There are likely multiple factors underlying the observed metadata decay. First, it is not surprising that older metadata are less likely to have been archived. Metadata archival practices are gradually improving, with more metadata being recorded into the INSDC, research papers, and online repositories such as Data Dryad (Figure 4A). This is consistent with increasing acknowledgement that these metadata are relevant and important to future research. However, the rate of metadata archival is apparently not keeping up with the rapid growth of genomic datasets (see Figure 1 of Toczydlowski et al., 2021) and it is certainly not closing the gap. Second, we found that authors of recent SRA datasets were significantly less likely to reply to our queries than those of older datasets (Figure 4B), although the overall response rate of 45% was comparable to previous studies (Vines et al., 2014, 2013). This result may indicate that recent SRA depositions are part of ongoing research projects for which authors are unwilling to share metadata for fear of getting “scooped” by others working on similar research questions. It is also true that younger authors are more likely to leave science than older authors (Reithmeier et al., 2019) and thus may no longer support their publications. Similarly, there may be a cohort effect in which authors of older studies are more established in their careers and have more time, and/or are more aware of increasing expectations around FAIR data, and thus more willing to communicate and share. Finally, of the authors that did reply, there was a significant decrease with the age of the BioProject in whether partial or complete spatiotemporal metadata were provided (Figure 4C,D), suggesting that if metadata are not properly archived to public repositories, they are subject to being lost over time, as previously highlighted for morphological data (Vines et al., 2014).

Taken together, our results support assertions that the current research system overly weights publications and citations, while underweighting scientific openness and transparency (S. W. Davies et al., 2021; McNutt et al., 2016; Nosek et al., 2015). Changing the system will likely require a combination of carrots and sticks (Whitlock, 2011). Carrots can take the form of citable data publications (Dimitrova et al., 2021), recognition of open data practices by hiring, promotion and tenure committees, or commendations from professional societies or departments (Roche, Kruuk, Lanfear, & Binning, 2015; Roche et al., 2014). Sticks in the form of open metadata mandates, must come from journals (Sibbett, Rieseberg, & Narum, 2020, Gareth Jenkins, pers. comm.), funding agencies, and data repositories, which all have a responsibility to respond to the needs of the research community (Lin et al., 2020). While we applaud the INSDC’s new spatiotemporal metadata annotation policy requiring country of origin metadata and their adoption of the MIxS metadata standards, we call for greater mandated spatial resolution to include at least a locality name or spatial coordinates (Table 1) with appropriate uncertainty or additional terms (such as Darwin Core’s coordinateUncertaintyInMeters, informationWithheld; Wieczorek et al., 2012) to protect endangered species or sovereignty of Indigenous Peoples (Hudson et al., 2020; McCartney et al., 2022).

Our datathon provided an unparalleled opportunity to train graduate students in the importance of proper data curation, and to raise awareness that almost every dataset has a potential for re-use. We suggest that training in data curation and metadata usage should be part of reproducible research training in every science graduate program, with emphasis on avoiding some of the metadata practices that hinder metadata recovery described in Box 1. Additionally, “datathons” such as that undertaken here could help to close the metadata gap in the short term, as they are very cost effective. If we assume a mean cost of sequencing of USD 50 per BioSample (and ignore the much higher, additional cost of sample collection and processing), this datathon rescued over USD 2.1 million worth of genomic sequence data for future research purposes. Co-authors of this paper spent about 2,300 hours on this metadata retrieval effort, which, if valued at an average wage of USD 19 per hour, yields a return on investment of nearly 4,700%, with average costs of remediating a BioSample or BioProject at USD 1.05 and USD 110 respectively. But ultimately, datathons are a stopgap solution.

Going forward, the entire biodiversity genomics research community should give the same priority to sharing metadata that they have given to sharing primary data, because it is only the metadata that make primary data FAIR. From a process standpoint, the collection of metadata should begin at the time of sampling, with the assignment of a globally unique identifier (GUID) to the actual material sample. This identifier, which should be assigned as early as possible after collection, serves as the root to which all subsequent derived products could be linked in an extended specimen cloud to establish clear provenance and thereby prevent duplication of data or effort (N. Davies et al., 2021; Lendemer et al., 2020). Through the use of GUIDs, both physical and digital products of the sample (digital sequence information, but also DNA or RNA extractions, subsamples, images, video, audio, CT scans, measurements of morphology, traits, gut contents, parasites, and other related data and associated metadata) will be linked to their material sample GUID to provide an extensive, holistic metadata cloud that can be used to better inform current research endeavors as well as create additional data-intensive research pathways. GEOME (Deck et al., 2017; Riginos et al., 2020) is an example of an easy-to-use “metadata broker” platform that can provide spreadsheet templates with definitions that can be filled in offline when the sample is collected. It can then mint a GUID for any sample that is added to it, and then harvest the INSDC accession numbers for genomic reads that are submitted to the SRA through GEOME, thereby maintaining permanent links between the sample metadata and genomic data.

The challenge then is to integrate these metadata downstream into databases (such as INSDC) which describe data derived from the sample. INSDC enables such linkages to other metadata platforms through the use of both Structured Voucher (https://www.ncbi.nlm.nih.gov/biocollections/docs/faq/) and Linkout (https://www.ncbi.nlm.nih.gov/projects/linkout/) facilities for both Nucleotide and SRA (through their corresponding BioSample record) datasets respectively (e.g. https://www.ncbi.nlm.nih.gov/nuccore/KC825472). Through these linkages, metadata corresponding to the original material sample can be tied to the resulting sequence(s) to both validate the metadata associated with the sequence record as well as provide updated information should specimens be reidentified or georeferenced after the lodging of the sequence with INSDC. Using the INSDC as a long-term repository for metadata about the sample may not make sense, in part because researchers who submit the sequences to INSDC have sole editing rights to the sequence record and it is currently quite difficult for others (such as the collections who hold the vouchers) to keep the INSDC metadata up to date or add additional information. Thus, the integration of these metadata from an upstream source somewhat negates the necessity for this information to be duplicated by the sequence depositor and ensures that the metadata are constantly up to date. This not only supports open, reproducible science (Buckner, Sanders, Faircloth, & Chakrabarty, 2021) but also exemplifies the Findable and Accessible principles of FAIR data (Wilkinson et al., 2016).

What this piecemeal system currently lacks, however, is support for data Interoperability and Reusability. This is because of the siloed nature of the data and our inability to compile it into a single resource for machine readability, data manipulation or downstream use. This shortcoming is being addressed through various initiatives such as the Extended Specimen Network (ESN; Lendemer et al., 2020; Thiers et al., 2021), the Digital Extended Specimen (https://dissco.tech/2020/03/31/what-is-a-digital-specimen/), the Distributed System for Scientific Collections (DiSCCo; https://www.dissco.eu/), iSamples (N. Davies et al., 2021) and others. Such a system would require all actors in the data landscape (researchers, collections, data aggregators, publishers, etc.) to utilize and publish resolvable GUIDs on all specimens, datasets and products of research to make these linkages possible, and thereby create an extensive online network of knowledge, and increase the potential for scientific research questions to be answered.

We join others in calling for ambitious policy goals that safeguard genetic diversity and scientific practices to enable this (Des Roches et al., 2021; Díaz et al., 2020; Laikre et al., 2020). Swift collective action is required to protect all levels of global biodiversity, and the first step towards protecting the evolutionary health of eukaryotic species worldwide is to close the metadata gap highlighted here. Simultaneously, conservation geneticists, molecular ecologists and evolutionary biologists must engage with global biodiversity assessment programs to ensure genomic data can be collected, interpreted and archived appropriately (Brodersen & Seehausen, 2014). Several exemplary international networks (e.g. GEOBON Genetic Composition Working Group, IUCN Conservation Genetics Specialist Group, and EU COST Action Genetic Biodiversity Knowledge for Ecosystem resilience [GBiKE]) have already made a case for protecting the genetic diversity of all species (Laikre et al., 2020), proposed indicators to gauge progress toward goals (Hoban et al., 2020; Laikre et al., 2020). These groups have asserted their rationale for these changes to stakeholders in policy documents, providing essential clarity in the use of genetic data, and reporting against targets (Hoban et al., 2021). These actions and advances encourage the uptake of genetic diversity monitoring by national authorities and international bodies. The vision for many of these biodiversity monitoring networks is to develop agile pipelines that intake raw biodiversity data and produce outputs that can directly inform conservation policies and decisions (Hoban et al., 2021). Yet, without appropriate archival of genomic data that includes the spatiotemporal metadata, we will be unable to deliver on the promise of genetic diversity monitoring.

The GEOME datathon enabled 13 students and 7 academics from 15 institutions and 4 countries to take account of the growing metadata gap for genomics data and begin to remediate it. The serendipity of being able to run a remote, distributed datathon due to travel restrictions and funding reallocation forced by COVID-19, in a time when Indigenous rights, biodiversity conservation and the value of genetic diversity have been front of mind, has not been lost on the participants. While our efforts have just begun to address the growing metadata gap, it is our hope that most researchers will start to ensure the FAIRness of their genomic data and metadata before or upon publication, thereby honoring the work that went into creating it and providing limitless opportunities for reuse of their data to help answer the important scientific questions of the future.

## Acknowledgements

This effort arose from an Evolution in Changing Seas Research Coordination Network (RCN) working group (NSF-OCE-1764316, Katie Lotterhos) and was funded by the Diversity of the Indo-Pacific Network RCN (NSF-DEB-1457848, to R.J.T.). We gratefully thank all of the authors who took the time to provide helpful responses to our metadata inquiries. We also thank Neil Davies, Chris Meyer, Beth Davis and Kiersey Nielsen for their input.

## Data Availability Statement

The metadata that were recovered by the datathon are available from the Genomic Observatories Metadatabase (GEOME; geome-db.org) https://geome-db.org/workbench/project-overview?projectId=305. Relevant code for analyses in this paper may be found at https://github.com/ericcrandall/geome_metadatathon1. The meta-metadata for BioProjects that were determined to be relevant to the datathon are in Supplemental Materials S4.

---

**Box 1.**
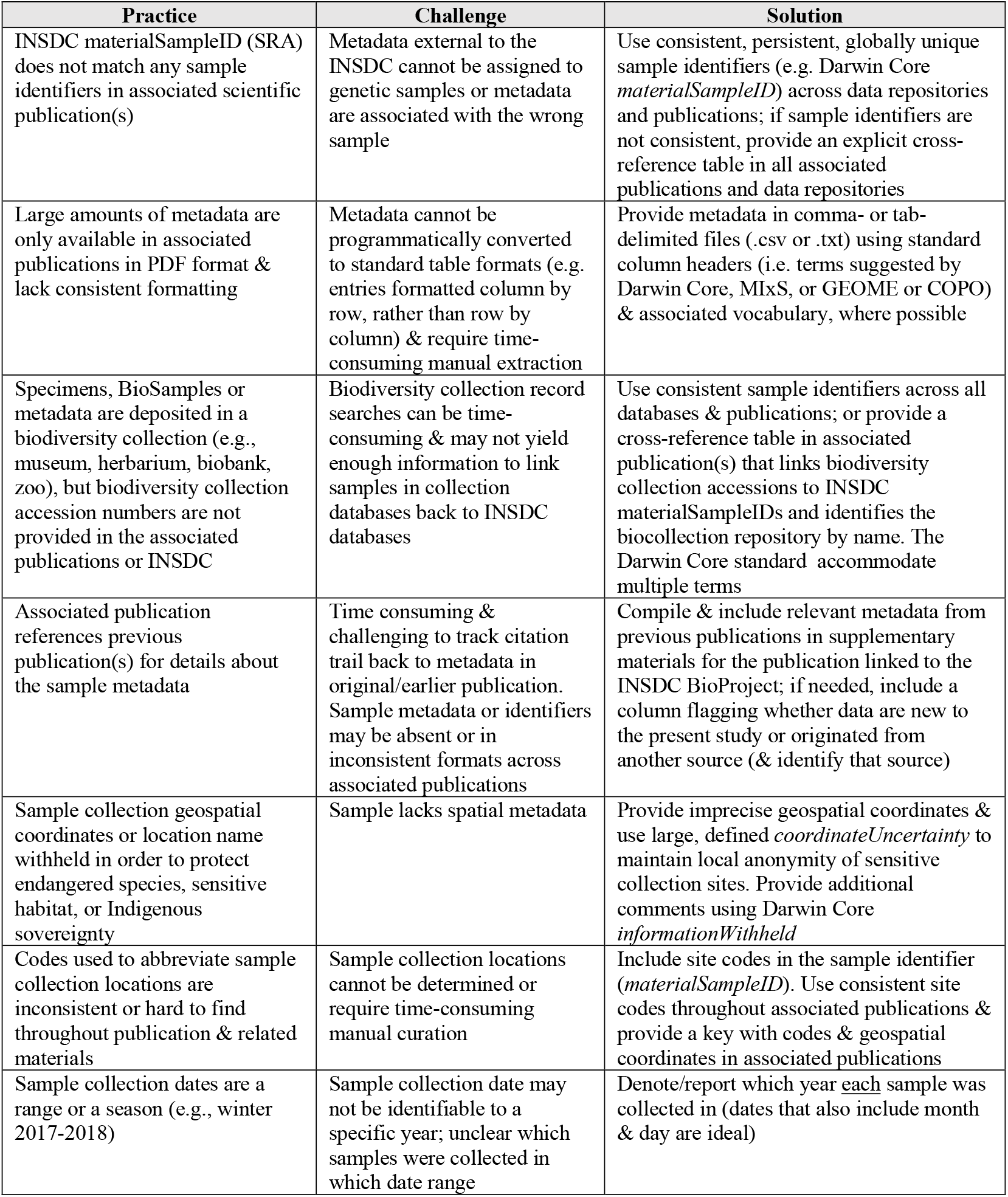
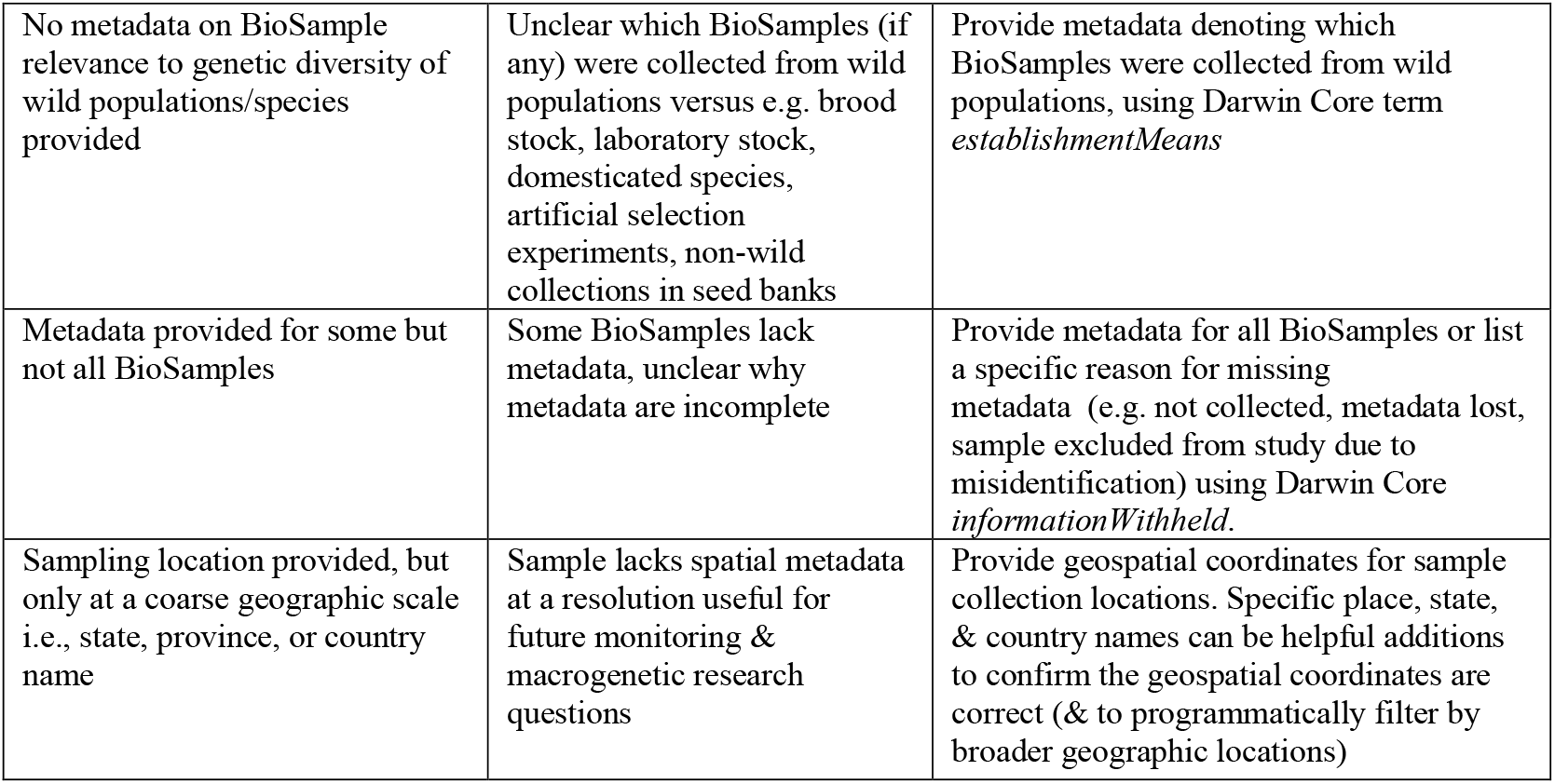
Summary of metadata practices encountered during the datathon that hindered metadata recovery and recommended practices to improve future usability of samples and genetic sequence data. In general, we recommend that authors use metadata software (GEOME: geome-db.org, COPO: copo-project.org, or museum database software such as Specify) to organize and archive their sample metadata.

