## Supplemental Materials S3 for "Metadata preservation and stewardship for genomic data is possible, but must happen now"

### Genomic Observatories Metadatabase (GEOME)
### Metadata Curation Protocol

Adapted 7/5/20 by Eric Crandall from protocols first developed by the Evolving Seas/DivDiv project.
Modified 7/23/20 by Eric Crandall to include structured comments about metadata provenance.
Modified 11/10/20 by Eric Crandall to add a section on quality control

#### Background:

Only about 20% of potentially biodiversity-relevant genetic data in the sequence read archive (SRA) of the International Nucleotide Sequence Collaboration (NCBI + DDBJ + EMBL) currently have geospatial metadata attached to them, meaning that the other 80% are nearly useless for meta-analysis of essential biodiversity variables such as heterozygosity, divergence and effective population size. The Genomics Observatories Metadatabase (GEOME) was created as a way to bridge the metadata gap between robust databasing efforts at the species and ecosystems level (GBIF, OBIS, BOLD) and at the genomic level (INSDC SRA). Please watch this 10-minute presentation for further backstory.

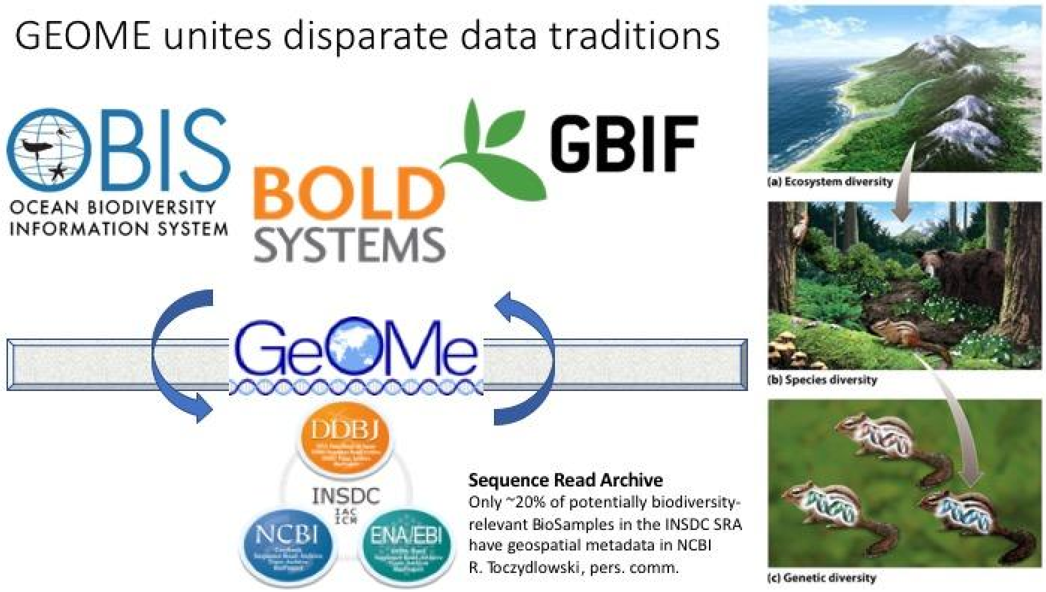


We downloaded metadata describing BioProjects containing reads from massively-parallel sequencing projects from SRA. The sequence data are collected in runs that have unique IDs like SRR5556902, SRR5556906, SRR5556101 that NCBI assigns, while each sample (which can have multiple runs) has unique IDs like SAMN1090410. In addition to the SRA ID that each run is labeled with (e.g. SRR5556906), many also have sample names that the authors provided (e.g. sample3_LakeSuperior). When the authors submitted these data to SRA, they did not provide the latitude and longitude of where each sample was collected. Authors *do* often report this geographic information in associated published scientific papers and supplemental materials. We need to find and link latitude and longitude (e.g. in a table in a published scientific paper) to each genetic sequence file that we have downloaded from SRA. Your challenge is to see if you can link the sample names that the authors provided to latitude and longitude information in published scientific papers or associated supplemental materials. The genetic sequence runs are grouped by sample, and then samples are grouped by datasets called BioProjects. Each BioProject often represents a unique study and a published scientific paper, so you will work to find metadata that describe BioProjects, which contain multiple samples and runs.

#### Project Communication

**Important Meetings And Dates:**

All Zoom Meetings to occur at: **https://psu.zoom.us/j/5664161782**

July 7th, 2020: Kickoff Meeting, 3PM EDT

July 8th, 2020: First Check-in Meeting at 4PM EDT. Meet every Wednesday thereafter at 4PM until August 21st.

August 21st, 2020: Final Meeting. All assigned BioProjects, or effort totalling ~100 hours must be completed by this date.

Outside of Zoom meetings, all communication about this project will occur via the #datathon Slack channel in the GEOME workspace: https://geome-workspace.slack.com/archives/C0164GWMDKR

If necessary, you can also contact Eric Crandall via

#### Miscellaneous Notes

Please see the Frequently Asked Question Document if you have a question that is not answered in this protocol. You may also consult or search the Slack #datathon channel.

You will probably not be able to work on a single BioProject and push it all the way to completion before moving on to the next one. In particular, there will be a necessary pause if authors of the dataset need to be contacted. Thus, you will likely want to be working on several BioProjects at the same time.

If you find multiple values that can go in a particular field (e.g. multiple DOIs for publications to go in the associatedReferences field), always separate these values with a pipe | (usually the capitalization above the \ key). **However**, when you have multiple values for any of the derivedData fields, please follow these instructions.

#### Metadata Protocol Overview

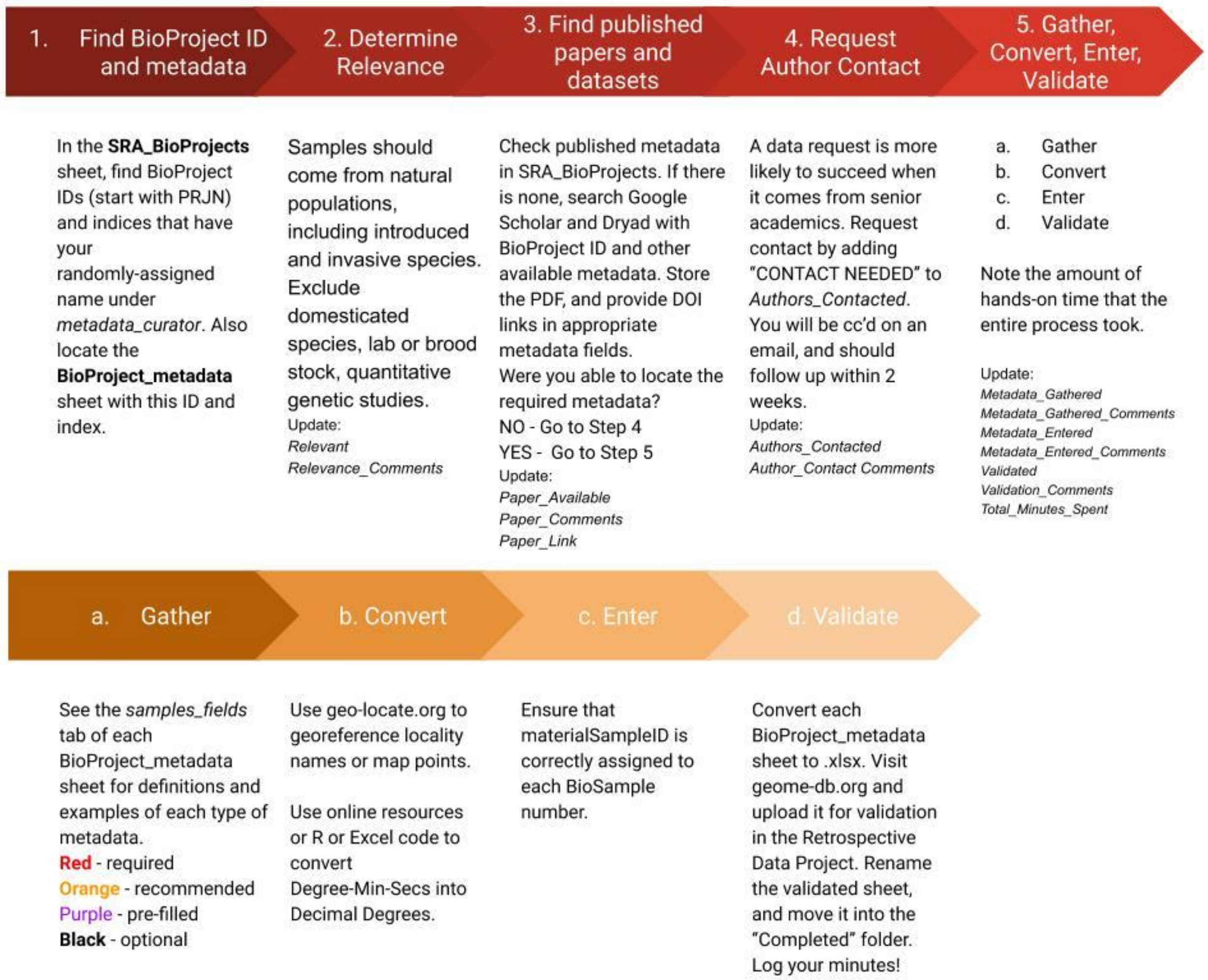


#### Complete Metadata Protocol

1. **Locate a BioProject ID** (starts with PRJN): in the first column of the **SRA_BioProjects** sheet to which your name has been randomly assigned (under *Metadata_Curator*). There will also be a corresponding BioProject_Index that will be easier to remember.

a. These SRA_BioProjects sheets (one for marine projects, one for non-marine projects) will be used to manage the datathon, and will be your starting place and where you will note your progress on each BioProject.
b. You should also be able to locate the corresponding **BioSample_Metadata** sheet in a subfolder.

2. **Determine Relevance to Genetic Biodiversity**: Try to determine the relevance of the BioProject for genetic biodiversity measurements from the metadata fields provided *project_title_bioprj* and *project_description_bioprj* should be especially useful here.
   a. Samples should come from natural populations, including introduced and invasive species.
   b. Pooled Seq is welcome, but we may have to think a bit about how to treat the metadata - contact an instructor.
   c. Studies have been filtered to those with at least 5 individual samples. If you encounter samples in a paper that do not appear on your metadata sheet, it is likely a species that is represented by fewer than 5 samples. Do not manually add these samples, but note this in *Paper_Comments*.
   d. In general exclude:
      i. Domesticated species
      ii. Broodstock, Laboratory Stock
      iii. Hybridization and quantitative genetics studies (wild hybrids are fine)
   e. If a BioProject contains a mixture of natural and captive samples, you can determine relevance if 50% or more of the BioSamples OR 50% or more of sampled populations are natural. You may flag the captive samples using the *establishmentMeans* field.
   f. You may need to locate and/or refer to associated published scientific paper/s (see #3 below) if you cannot confidently determine the relevance of the dataset using the BioProjects sheet.
   g. If in doubt, ask an instructor!
   h. Write your determination of relevance under *Relevant* as a Boolean value (TRUE or FALSE). Give a brief explanation of why you determined the dataset was or was not using the controlled vocabulary under *Relevance_Comments*.

3. **Find Associated Published Papers**: If nobody has found a paper for this BioProject yet (*Paper_Available* = NA or Blank, and there is no link under *Paper_Link*) then try to find scientific publication(s) derived from this dataset:
    a. The first place to check is the metadata fields starting with *project_target_scope* all the way to the end on this same sheet. Some (but not most) BioProjects have information about the associated publication in these fields.
    b. Second, try searching for the PRJN number. Searching directly for the BioProject ID number *(project_acc_bioprj*) in Google Scholar is sometimes fruitful, as authors sometimes list this in a data availability statement in an associated published paper. See Rachel's tips about locating the correct paper here.
    c. If neither steps a or b work, try looking at at *first_name_biosample* and *last_name_biosample*, or *contact_name_sra_all*, to find the name of the person and/or institution that submitted the data to the SRA. Then look at *project_name_bioprj*, *project_title_bioprj* and *project_description_bioprj*. Try pasting these values, together with the organism's common or scientific names into Google Scholar, together with the responsible party's name, and see what comes up.
    d. You may have to search on several iterations of these and other metadata fields until you find a likely paper. You should be able to judge whether you have found the right paper from the title and abstract of the paper.
    e. An informative paper will have one or more of:
        i. A map of sample sites
        ii. A table of sample sites with lat/longs
        iii. A table of sample sites with lat/longs in the supplemental online information.
        iv. Sample site names given directly in the methods (likely under a sample collection type heading).
        v. Your coordinateUncertainty in meters should be less than 100,000 (100 km). If you can't narrow down the sampling locality to less than 100 km, you should request to contact the authors.
    f. If you find an informative paper:
        i. Change the *Paper_Available* field of the **SRA_BioProjects** sheet to TRUE.
        ii. Put the full DOI (including http://doi.org/) for the paper in the *Paper_Link* field. State exactly where in the paper you found the lat/long information (Table or Figure #) in *Paper_Comments*. If you have multiple papers, separate the DOIs with |

iii. You will also need to add this | separated list of DOIs in the *associatedReferences* field of the **BioProject_metadata** sheet for this project.
iv. Download the PDF, renaming the file with BioProjectID_firstAuthorLastName_year.pdf. Place the PDF in the shared google drive under the Papers directory.
v. If there is pertinent supplemental data, rename it following the name naming convention above, but add _SuppInfo to the filename. Place it in the same folder.
vi. Go to Step 5 in this protocol to fill in the metadata.

g. If you cannot find an informative paper within ~30-45 minutes of trying, then write FALSE under *Paper_Available*. Go to Step 4 in this protocol.

4. **Request Author Contact**: If you cannot find a paper with informative metadata about the samples, or if the metadata for **required** fields are incomplete, the authors of the dataset will need to be contacted. Be sure that you have checked through all supplemental or DRYAD or other online data sources mentioned in the paper before you request contact. In our experience, it is often helpful to have someone at a professor level make the initial contact, and they may know the authors or be acquainted through their personal network. Progress on this dataset will need to be paused until the following steps are completed:
    a. First, fill in the metadata sheet with all metadata that you have been able to find online. This will help to demonstrate that we have made an attempt.
    b. Add "CONTACT NEEDED" to the *Authors_Contacted* field. This will alert GEOME personnel to the need to make an initial contact.
    c. Add any specifics about what data are missing, or why the authors need to be contacted to *Author_Contact_Comments.* Add these comments in a structured format: Copy and paste the following into the Author_Contact_Comments, and add "TRUE" after metadata fields for which you have data for every sample, "MOST" if you have metadata for >50%, "SOME" if you have metadata for < 50%, and "Reason" (incl. quotes) after fields for which there is no metadata, where reasons can be: "There is no paper associated with this project" or "Table 2 in Wallace et al. only includes states as localities, uncertainty > 100km", etc.

```
materialSampleID =
locality =
coordinates =
country =
habitat =
```

environmentalMedium =
yearCollected =
permitInformation =
preservative =
derivedGeneticDataX =

d. Items in these fields should be delimited with |
e. If you are missing data for all fields, you may simply state all = “Reason”
f. If you have partial data for certain fields, substitute “MOST” for “TRUE” if more than 50% of samples have metadata and “SOME” if less than 50% have metadata.
g. Move on to other BioProjects.
h. GEOME instructors will try to find someone with a personal connection to the dataset owners. They will then send a form-letter email from, co-signed by several instructors and you, with your email address in cc. The email will contain the Bioproject_metadata sheet for them to fill out. The instructor will add their name under *Who_Will_Contact* and add the date of their email under *Email_Date*.
i. It will also be your job to follow up with the author if you have not heard back from them in two weeks.
j. Once we receive a reply with the requested metadata, it will be up to you to add a date under *Author_Response_Date*, and fill out *Author_Response_Comments* with the same structured comment format as above. If you already had metadata (e.g. for locality) from the paper, your comment for this field will **remain** “TRUE”. Generally, you can just copy and paste *Author_Contact_Comments* into *Author_Response_Comments*, changing fields that you received to “TRUE”. If you received no response from the author, then you can just copy and paste the text, leaving it unchanged.
k. If you are not able to obtain the required metadata from either a publication or from the author, note FALSE under *Metadata_Gathered*, and describe the circumstances under *Metadata_Gatherered_Comments*. Be sure to still fill in any metadata that you *do* have.

5. **Gather, Convert, Enter and Validate Metadata**: If you’ve obtained metadata from a publication or from the author themselves, nice work! Add TRUE to the **SRA_BioProject** field *Metadata_Gathered*. For *Metadata_Gathered_Comments* write either “From Author” or “From Paper” or “From DRYAD or elsewhere” (exactly these phrases). You can now move to  the BioSample_metadata table,

located in /BioProject_Tables/(Non)Marine_BioSample_Metadata, and named with the BioProject index and ID.

a. **Gather Metadata**

   i. You will be entering metadata in the *Samples* tab of the BioSample_metadata table (tabs are located across the bottom of the sheet). Detailed instructions are given on the *instructions* tab, and the *Samples_Fields* tab. Controlled vocabularies are given in the *Lists* tab. Controlled vocabularies mean you need to pick a specific term from a given list. It is very important that you stick to the controlled vocabularies because we will use these in coding and organizing all datasets later.

      1. The **Red** fields indicate fields that are required.
      2. **Orange** fields are recommended. Fill these in if at all possible.
      3. The **Light Purple** fields should all have been pre-filled with metadata already in the SRA. However, there may be obvious errors or issues with this pre-filled metadata, or it may be missing.
      4. **Purple** fields should be pre-filled. Please don't change any information in them. If information is missing, please alert an instructor.
      5. **Black** fields are optional. The more complete you can make this metadata, the more useful it will be, but don't get bogged down with completing every single field! Also, don't guess at values!

   ii. You might want to use tabula (https://tabula.technology) to retrieve information from tables in PDFs

   iii. *materialSampleID* is an important field because it lists the original identifiers that the investigators gave to their samples. This might be the only thing that ties the actual physical sample to its genetic sequences. In many cases it should have been pre-filled from SRA metadata, but if it isn't you should take a look in the *library_name* field of the main BioProject sheet. This may have a list of library names, which may or may not be equivalent to sample names. Be sure to find evidence in the paper or supp. Data that the library names are relevant to the sample identifiers.

   iv. *establishmentMeans* is another important field for helping future investigators determine how "natural" a sample is. If your sample comes from the "wild" it could be "native", "introduced", or "invasive". If it comes from captivity, it might be one of several categories of experiment, managed (as in domesticated, agricultural, seed bank etc.) or lab stock.

v. You might find information about the sampling permit (*permitInformation*) that was used to obtain the samples in the acknowledgements or somewhere following the Discussion. Minimally, we'd want the permit number and issuing authority. Permits may differ by locality, especially if samples were taken in different states or countries. This is important info if you can get it!

vi. We would like to have all associated publications (as complete DOIs) placed in *associatedReferences*. You can just copy the *Paper_link* field from the master sheet (DOIs separated by |). If you can't find a DOI, then a standard URL will work too.

vii. Be careful when diagnosing the habitat for your samples. If it's clear that a sample came from the marine benthos, then you can pick *marine benthic biome*. If it isn't clear, then maybe *marine biome* would be better. Also, be aware that habitat can change at the level of each individual sample. For example some individuals in a project might have been sampled in an estuary and others might have been sampled in pelagic ocean. Encyclopedia of Life can be an authoritative source for habitat (search your species, click data tab, filter attributes for habitat). As always, direct any questions to datathon instructors.

viii. Check the "Data Accessibility" or similar section at the end of the paper for derived datasets. Derived data can be defined as any dataset that was produced from the raw data following processing through some analytical pipeline that involved some subjective decisions by the user. Such data include SNP calls, microsatellite data, and OTU or ASV tables. These data are often stored in a repository like Data Dryad, and it should be sufficient to add the link to the dataset, and the filename, as well as what type of derived data it is, and the format of the file.

    1. **Important Note!** When there are multiple values for most fields, we've asked you to use | to separate them. However, when there are multiple derived datasets for a sample, you need to:
        a. First fill in all other metadata
        b. Highlight and copy all metadata rows, pasting them below the first group of entries.

c. Change ONLY the derived dataset fields to point to the second derived dataset.
d. Continue this process of copying rows and changing derived dataset fields for every derived dataset that you link to.

**b. Convert Metadata**

i. Verify that latitude and longitude data are in decimal degrees (DD), not degrees-minutes-seconds (DMS). For latitude, a negative value indicates southern hemisphere, whereas for longitude, a negative value indicates western hemisphere.
   1. Coordinates for Grand Prismatic Spring, Yellowstone NP
      a. DMS: N44° 31' 30.1764" W110° 50' 17.4834"
      b. DD (we want this): 44.525049, -110.83819

ii. Non-decimal-degrees may be converted [here](http://data.canadensys.net/tools/coordinates), or in R or Excel. http://data.canadensys.net/tools/coordinates

iii. If all you have is a map or a list of place names, use GeoLocate ([https://www.geo-locate.org](https://www.geo-locate.org)) to georeference these, with appropriate uncertainty noted in *coordinateUncertaintyInMeters* field. Coordinate uncertainty should describe the radius of a circle within which the sample was or could have been taken. Thus, if the only locality information says that a sample came from "San Francisco Bay" the minimum coordinate uncertainty *in meters* is 68,000. "Edit Uncertainty" in GeoLocate will allow you to estimate this nicely.

**c. Enter Metadata**

i. When entering the metadata, you'll find that there is a lot of repetition in that same value sometimes has to be entered for every BioSample and Run. FYI, GEOME is a relational database, so for example values that are common to a particular sampling event at a particular locality will be kept in an "Events" table, thereby reducing redundancy. But for now, enter the redundant values.

ii. If your project number is less than E0423 or M058 then you will need to change voucherURI field to establishmentMeans, and refer to the controlled vocabulary on the [project template](project template).

iii. Consider sorting the metadata table so that all samples from a particular locality group together. This will make it easy to click'n'drag the same values into the right places. **If you sort, please sort the entire sheet.**

iv. **Take extra care** that the identifiers originally given by the BioProject authors (in materialSampleID) line up correctly with their BioSample IDs. If you have sorted as in step ii above then the values found in library_names in the main sheet will no longer correlate to the BioSample accession numbers. You can re-sort to the original order using *Project_Index.*

**d. Validate Metadata**

i. Download .xlsx spreadsheets for each of your completed **BioSample Metadata** Google sheets.

ii. Go to the GEOME website (geome-db.org). Click on "Workbench". Then choose "Retrospective SRA Datathon" from the drop-down list at the top of the screen. No need to login. Then from the menu on the left, select "Validate Data". Click the check boxes next to "Excel Workbook" and "Only Validate Data".

1. The interface should show you a map of where your samples are located. This should make sense with what you know from the study!
2. Move through all validation errors and warnings, doing your best to fix them, or consulting with an instructor.
3. You may ignore any warnings having to do with ExpeditionCode or Expedition-anything (no need to even paste them into the validation comments field).

iii. Once your metadata are successfully validated, write TRUE in *Validated* of the master BioProjects sheet and give any comments about warnings that couldn't be avoided etc. in *Validation_Comments.* If there are warnings that can't be avoided, copy and paste the entire text into *Validation_Comments.*

iv. Add "_validated" to the end of the spreadsheet filename, and place in the "Completed_..._BioSample_Metadata_Sheets" folder.

v. Great job! You've completed the metadata for this BioProject. Be sure to note the approximate number of hands-on minutes you spent on this BioProject in the total_minutes_spent column of the SRA_BioProjects sheet before moving on to your next assigned BioProject.

6. Quality Control (added November 2020, modified May 2021)
    a. Find projects that have been validated but not QC'd using the appropriate SRA_BioProjects sheet (marine or non-marine).
    b. After selecting a particular BioProject to QC, check that all other fields in the master non-Marine SRA BioProjects sheet make sense for that BioProject,
        i. Metadata_Gathered and Metadata_Entered are both TRUE
        ii. Author_Contact_Comments and Author_Response_Comments are in alignment.
        iii. Read through any notes left by the curators about outstanding issues with the dataset.
    c. All validated projects should have an .xlsx in the Completed_BioSample_Metadata_Sheets. Open this file in Excel
    d. Sort the sheet by locality. You should see that each distinct locality has a distinct set of coordinates. If it has been validated, then the student has already checked that these coordinates match those in the paper.
    e. Re-sort by sheet_index. Check that each metadata field contains reasonable values. Each red-coded field should be filled, and hopefully many orange fields. <u>If you can easily add missing information about habitat, establishmentMeans, environmentalMedium etc., do so</u> (noting this in Quality_Control_Comments).
    f. If there are missing values for locality AND decimalLatitude + decimalLongitude, then delete those entries entirely. Other missing information is not as critical, and we will upload these to GEOME anyway after consultation with John Deck.
    g. Re-sort the spreadsheet by sheet_index. DOIs in **associatedReferences** and **derivedGeneticDataURI** should almost always be uniform across the project. In many cases, the student has click'n'dragged such that these values increment. <u>You may need to fix this</u>. Also, be sure to add the associatedReferences DOI if it is not present (there have only been about 20 BioProjects that didn't have one thus far). Click and follow links in both fields to make sure they are correct.
    h. Carefully check all of the derivedGeneticDataX fields. There were a lot of misunderstandings among the data curators about what constitutes derivedGeneticData. We are only accepting live links to actual genetic data, not summary statistics in this field. Recall that all sample entries need to be copied for every new derivedGeneticData file that is added. If

there is a description of the file in the linked URL (mandatory in Dryad) include that in **derivedGeneticDataRemarks**.

i. Double-check the purple and black pre-filled and optional fields to the right side of the sheet.
j. Save the spreadsheet into the “QC” folder, appending “_QC” to the filename.
k. Re-check all the fields for this BioProject in the SRA_BioProjects master sheet, making sure they all reflect the final status of this BioProject.
l. Mark the project as QCd in the SRA_BioProjects sheet (change “FALSE” to “TRUE”), making any notes about changes you made.

**Metadata Checklist**

**Required**, **Recommended**, **Optional**

**Pre-Filled** (hopefully)

| |
|---|
| materialSampleID |
| decimalLatitude |
| decimalLongitude |
| coordinateUncertaintyInMeters |
| georeferenceProtocol |
| locality |
| habitat |
| environmentalMedium |
| sampleEnteredBy |
| yearCollected |
| country |
| establishmentMeans |
| derivedGeneticDataType |
| derivedGeneticDataURI |
| derivedGeneticDataFormat |
| derivedGeneticDataFilename |
| permitInformation |
| preservative |
| island |
| landOwner |
| lifeStage |
| maximumDepthInMeters |
| maximumElevationInMeters |
| microHabitat |
| minimumDepthInMeters |
| minimumElevationInMeters |
| monthCollected |
| derivedGeneticDataRemarks |
| samplingProtocol |

| |
|---|
| sex |
| stateProvince |
| tissueType |
| voucherURI |
| continentOcean |
| dayCollected |
| otherCatalogNumbers |
| principalInvestigator |
| phylum |
| class |
| order |
| family |
| genus |
| specificEpithet |
| infraspecies |
| colloquialName |
| nomenclaturalCode |
| collectorList |
| project_acc_bioprj |
| run_acc_sra |
| biosample_acc_sra |
| sample_acc_sra |
| library_selection_sra |
| library_strat_sra |
| read_type_sra |
| sequencing_platform |
| library_prepraration_protocol |
| instrument_model |
| study_acc_sra |
| experiment_acc_sra |
| package_biosamp |

**dataset_id**
