## Supplemental Materials S5 for "Metadata preservation and stewardship for genomic data is possible, but must happen now"

Dear Drs. {FirstAuthorName},

I am emailing you regarding your {Year} study, cited below:

{References}

As you know, published data often have utility beyond the original publication. This is likely to be increasingly the case with data describing genetic diversity, which are so vital to basic science and conservation efforts.

We are affiliated with a collaborative effort to aggregate genetic data, which includes individuals  affiliated with the Genomic Observatories Metadatabase ([http://geome-db.org](http://geome-db.org/)), The Diversity of the Indo-Pacific Network ([http://diversityindopacific.net](http://diversityindopacific.net/)), and the Ira Moana - Genes of the Sea - Project (<https://sites.massey.ac.nz/iramoana/>) with funding from the National Science Foundation and the Royal Society of New Zealand Te Apārangi. Although these research projects are focused on marine biodiversity, we are presently compiling metadata for all of Earth’s ecosystems.

**We have retrieved metadata for genetic sequence data from the Sequence Read Archive from a publicly-available project for which you are listed as the contact author. Unfortunately, we were unable to locate the following key information in NCBI regarding where and/or when these genotyped individuals were collected:**

1) {Reason1}

2) {Reason2}

We realize that including such metadata within the SRA can be challenging. However, missing links between raw sequence data and metadata limit the long-term utility of your hard-won data to the wider field of molecular ecologists and conservation geneticists.

**Attached is a spreadsheet to fill in that will provide additional context for your published data. Red** fields indicate metadata that we view as necessary for basic analysis of genomic biodiversity. **Orange** fields are recommended, while **black** fields are optional. **Purple** fields and any other filled fields are based on our translation of information provided in your BioProject or publication and you should only need to confirm that they are correct. We ask that you fill this sheet in and return this as soon as possible or nominate a time by which you will be able to do this. We have done our best to populate this sheet with metadata that are available in your paper or elsewhere; please forgive us if we have overlooked something. You are welcome to provide any additional information in blank fields or refine or correct the information we have provided.

The information you return will be added to the Genomic Observatories Metadatabase ([http://geome-db.org](http://geome-db.org/)), which provides open-access permanent links between sample/collection metadata and SRA records. Please also see [our recent paper](https://onlinelibrary.wiley.com/doi/full/10.1111/1755-0998.13269) and [editorial endorsement](https://onlinelibrary.wiley.com/doi/10.1111/1755-0998.13283) in Molecular Ecology Resources. Thus, your collection data will be available to future researchers adding value to your publicly accessible genetic data already in the SRA. (Indeed, we encourage you to consider submitting your future genomic data through GEOME, which can facilitate preparation for NCBI submission).

**We look forward to your response as to whether you will be able to assist us and will follow up with you in two weeks if we have not yet heard back**. {GradStudent} is a graduate student who is assisting with this effort, please cc them in your response. We have aggregated a FAQ of common questions that you can refer to [here](https://docs.google.com/document/d/1IFTHb5BKhuA6ZkQYSyNdpY4RCmDtBFtak_d5fyOhAFY/edit). If you have any additional questions or are not the correct contact author, please let us know as soon as possible.

Thank you for your help in preserving the long-term utility and integrity of genomic data!

Sincerely,

{GradStudentHonorific} {GradStudent} – {GradStudentInstitution}

Dr. Eric Crandall - Pennsylvania State University

Dr. Michelle Gaither - University of Central Florida

Dr. Libby Liggins - Massey University

Dr. Cory Noble - Massey University

Dr. Rachel Toczydlowski - Michigan State University

Dr. Cynthia Riginos - University of Queensland
