## Supplemental Materials S6 for "Metadata preservation and stewardship for genomic data is possible, but must happen now"

### 1. Effect of BioProject Age on p(associatedReferences)

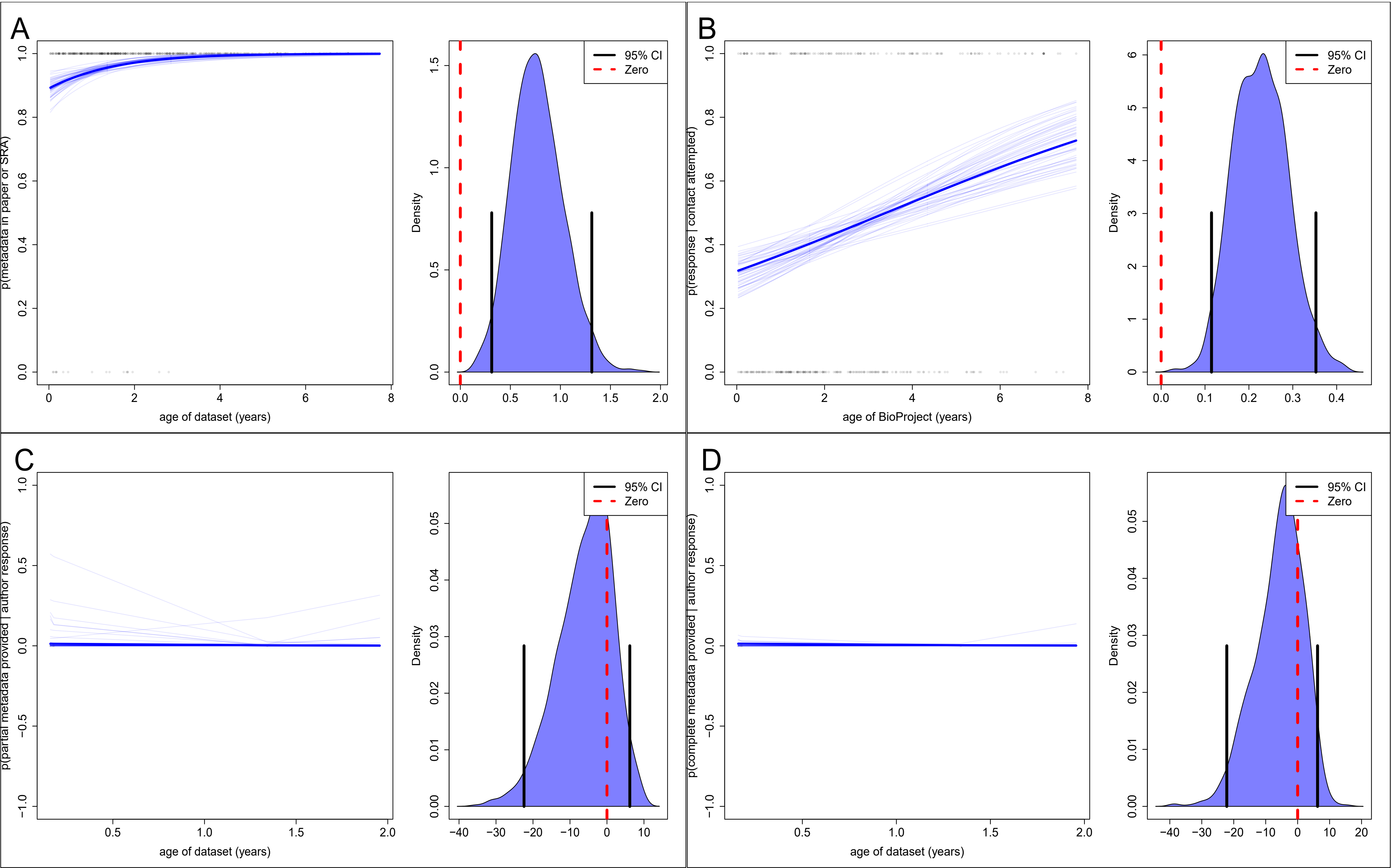

### 2. Effect of BioProject Age on p(decimalLat.Long)

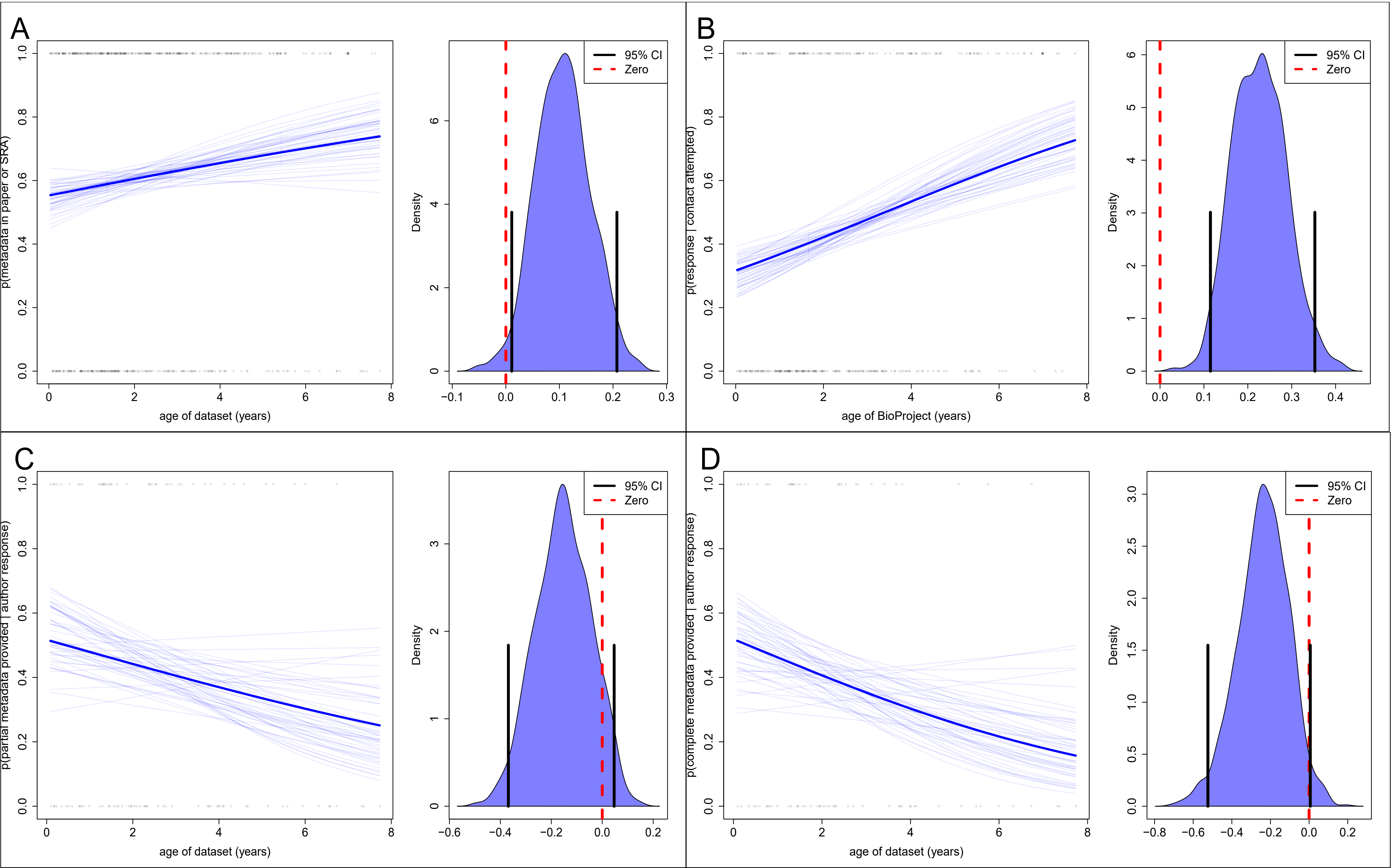

3. Effect of BioProject Age on p(country)

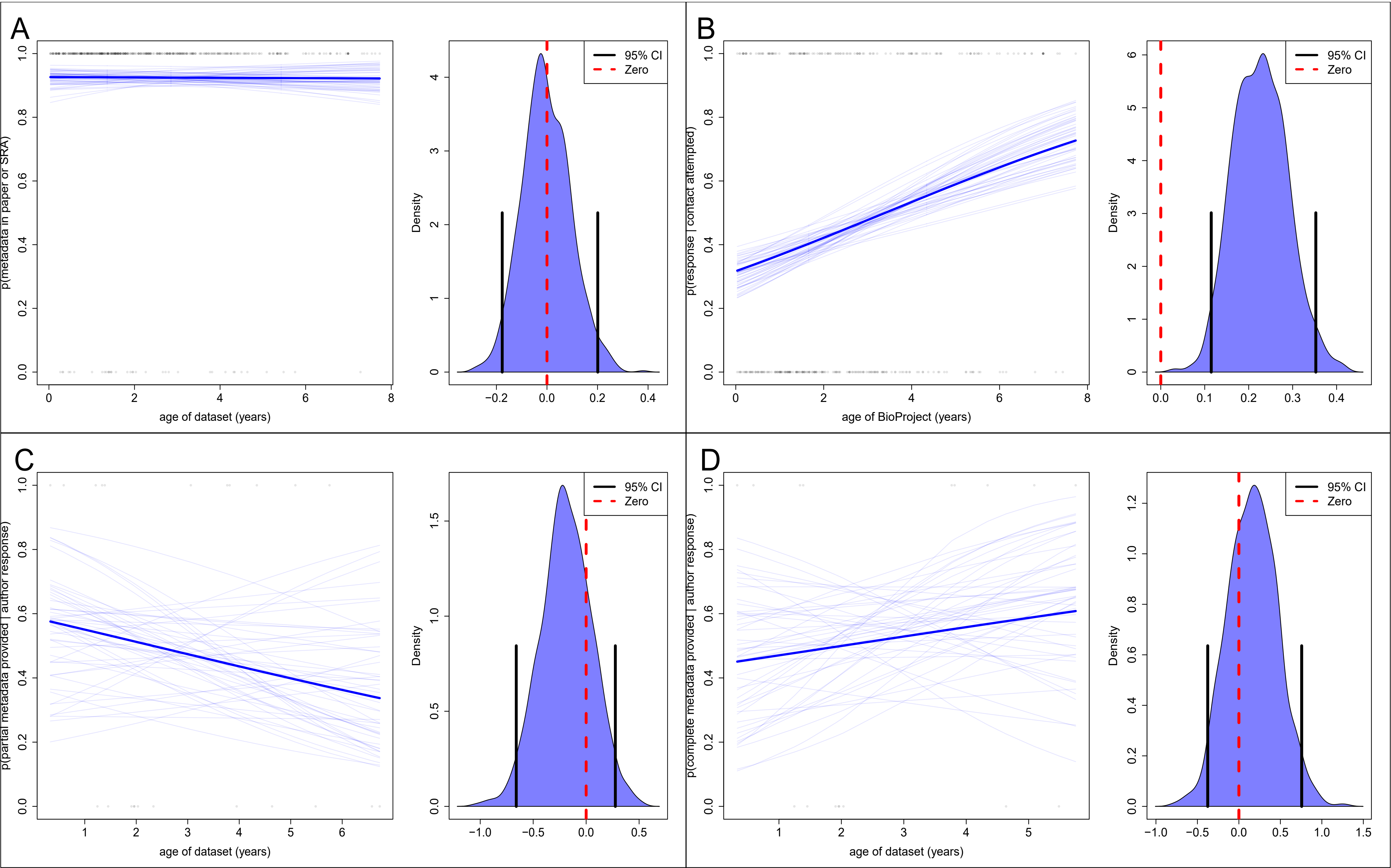

### 4. Effect of BioProject Age on p(derivedGeneticData)

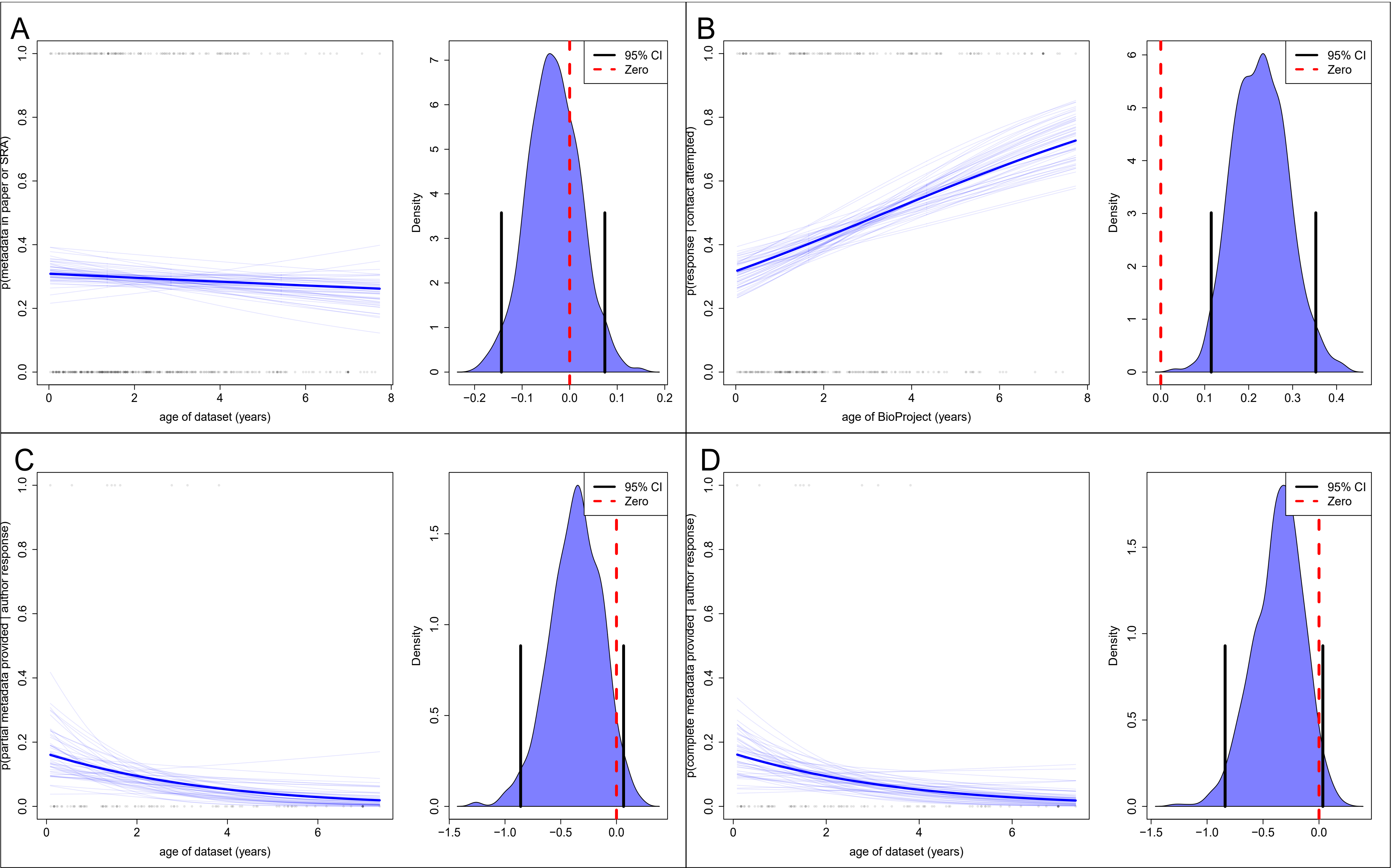

### 5 Effect of BioProject Age on p(environmental_medium)

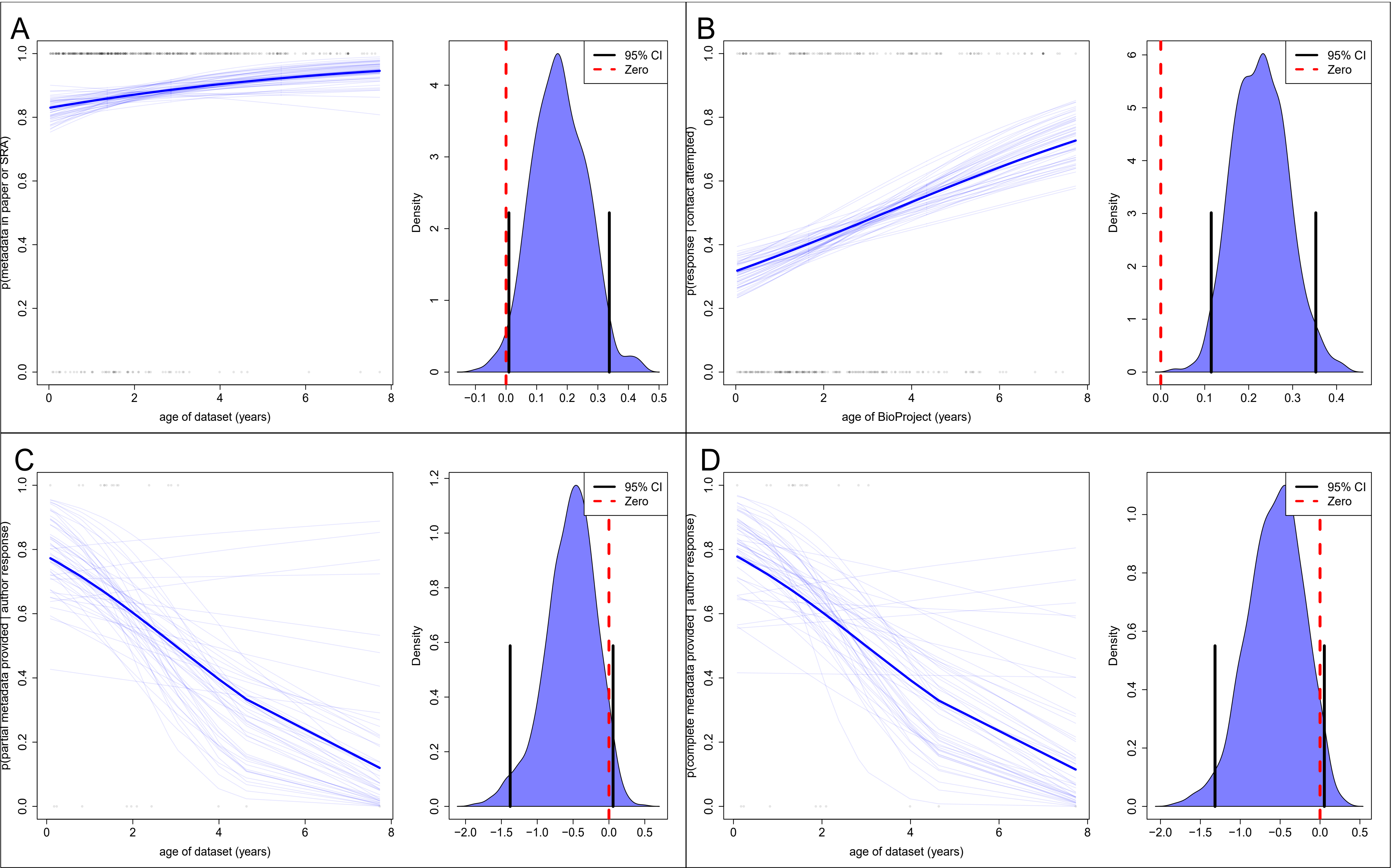

### 6. Effect of BioProject Age on p(habitat)

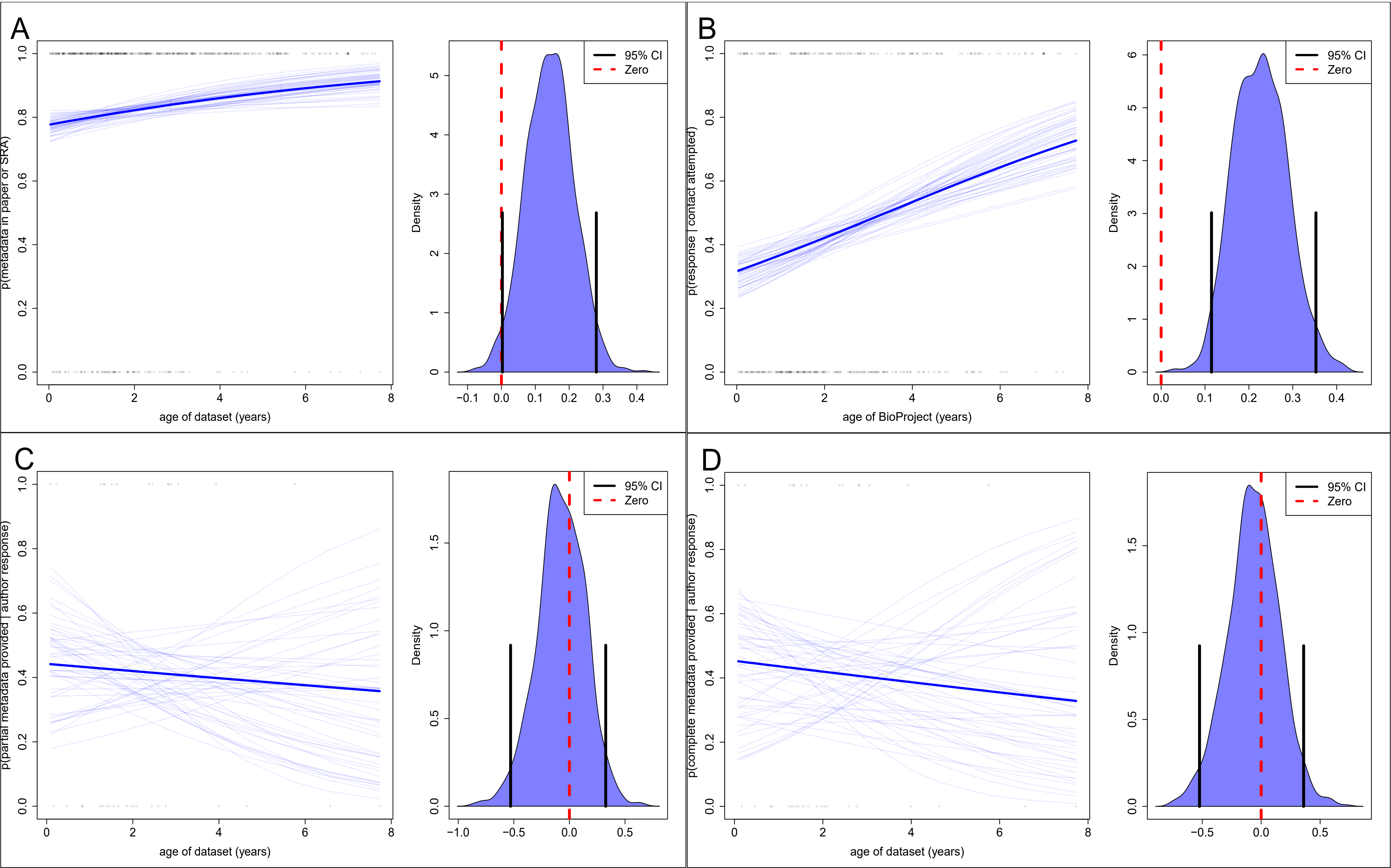

### 7. Effect of BioProject Age on p(locality)

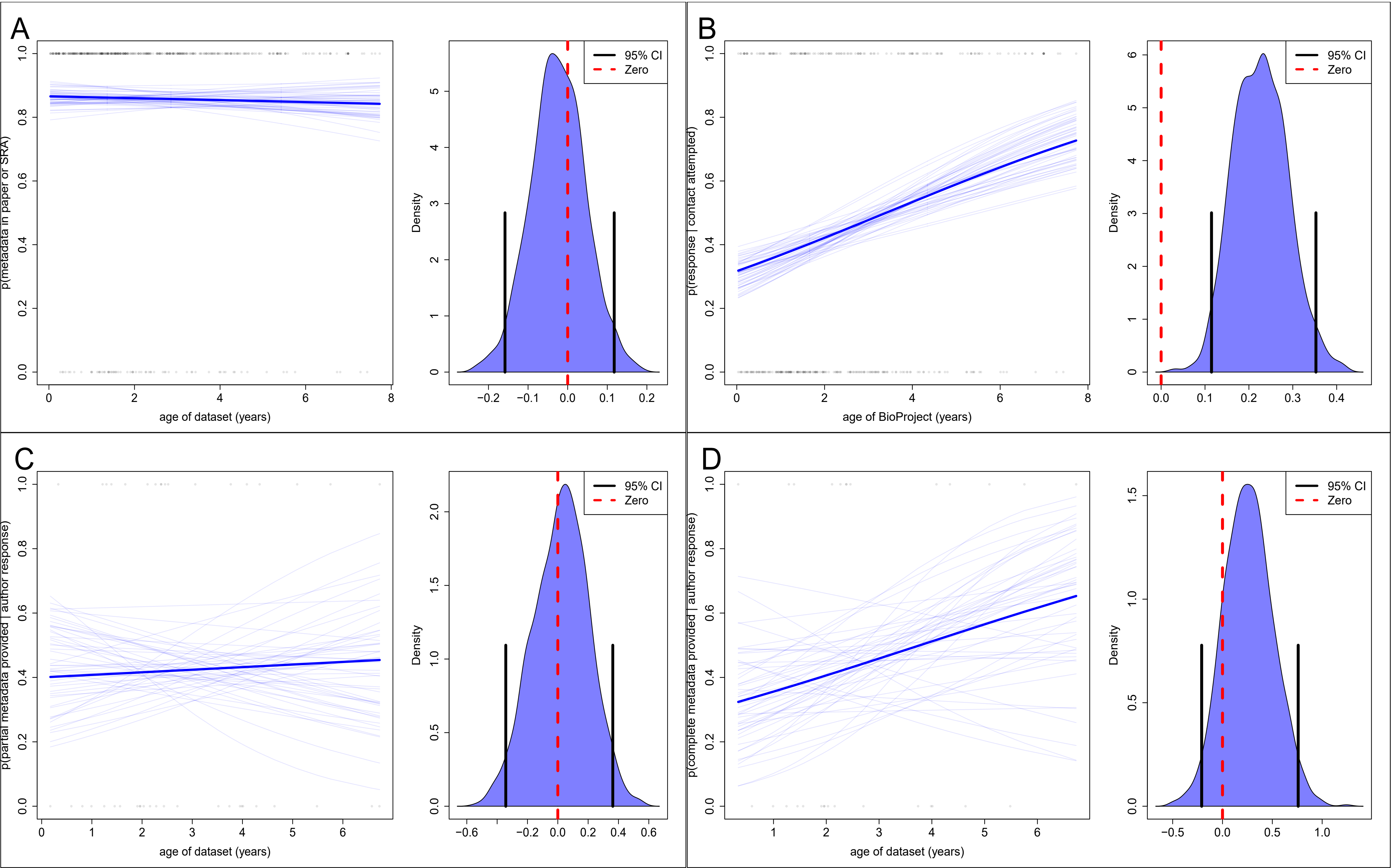

### 8. Effect of BioProject Age on p(permitInformation)

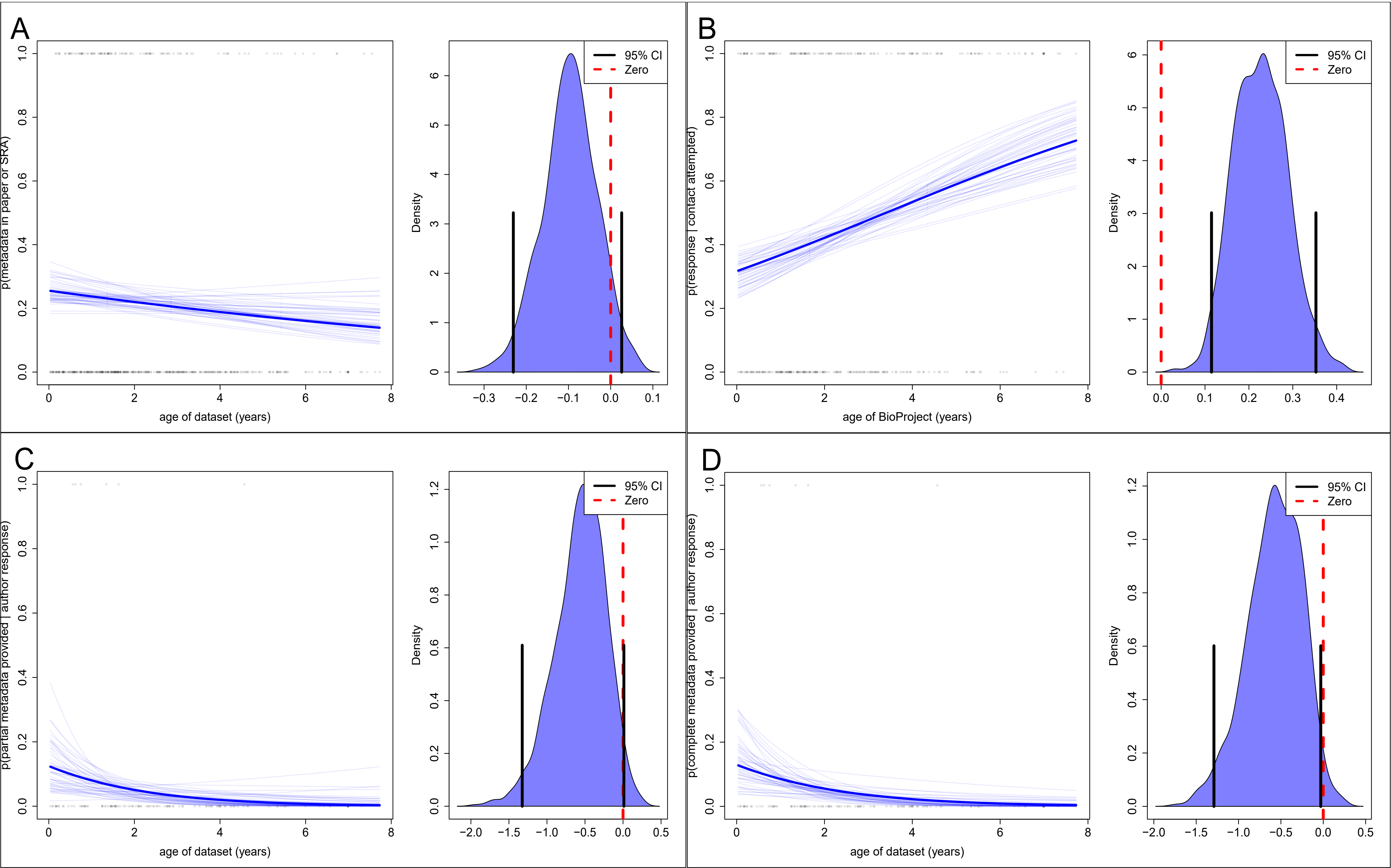

### 9. Effect of BioProject Age on p(preservative)

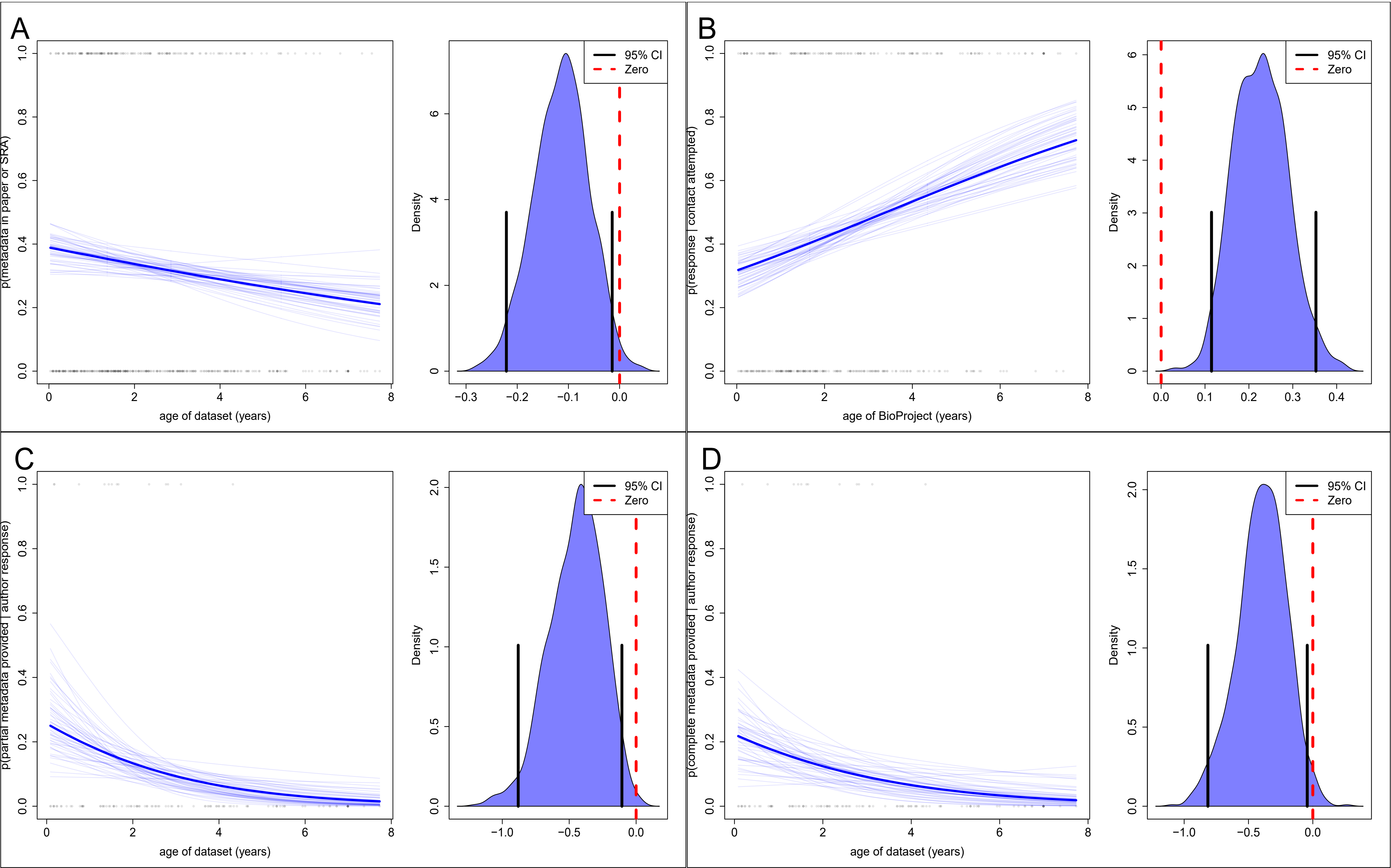

### 10. Effect of BioProject Age on p(materialSampleID)

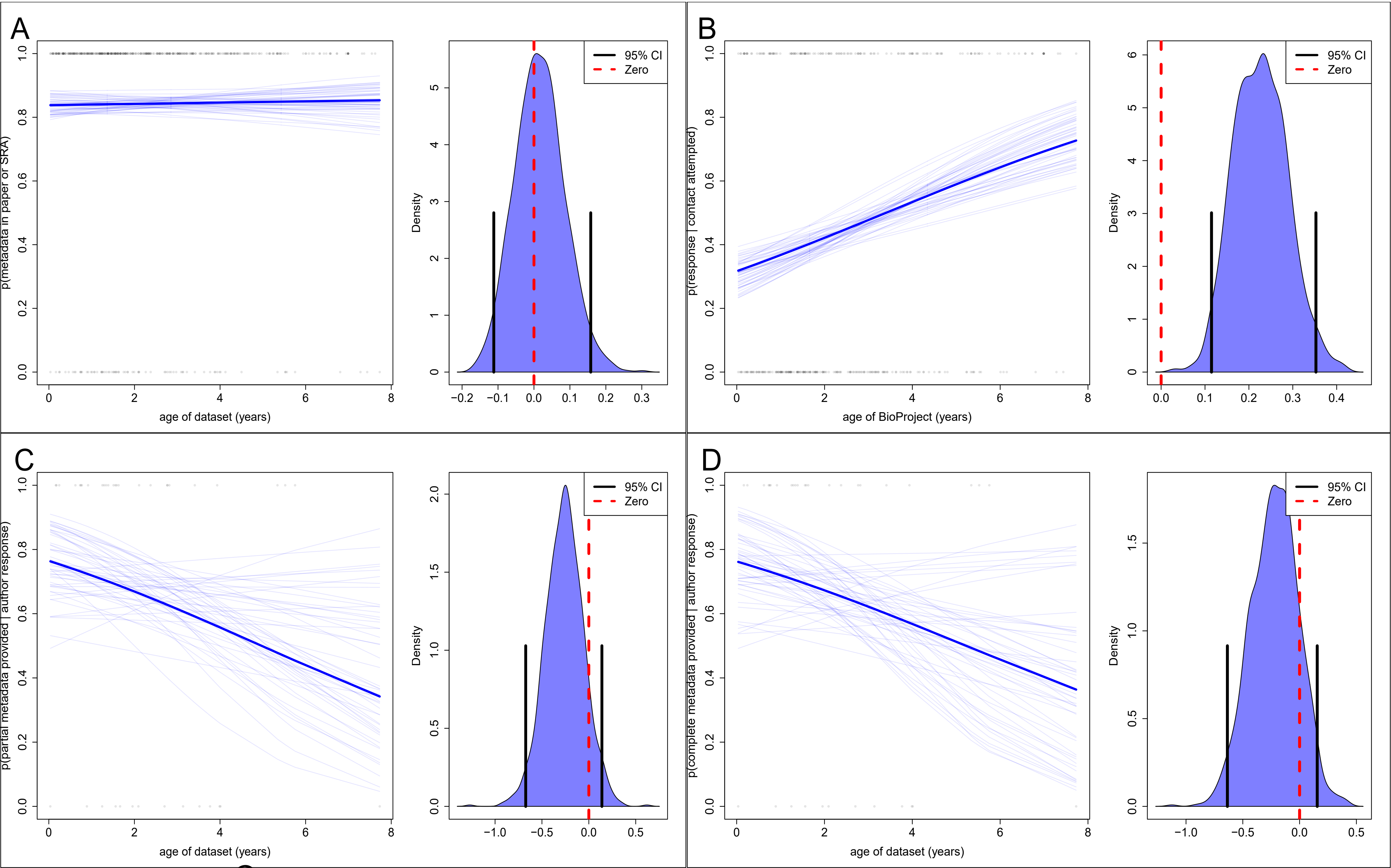

11. Effect of BioProject Age on p(yearCollected)

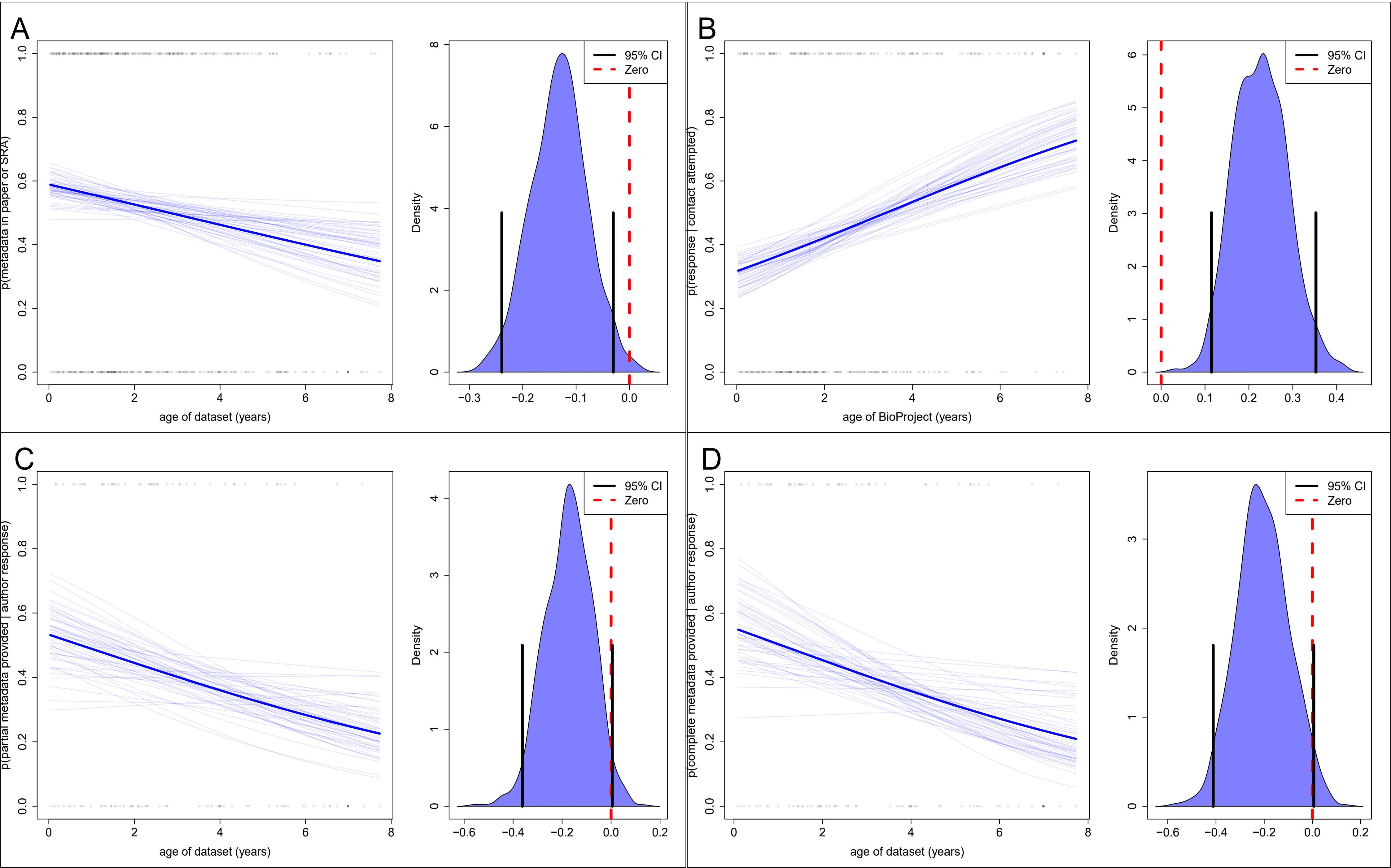
